## Supplementary Information for "Heterotypic interactions in the dilute phase can drive co-condensation of prion-like low-complexity domains of FET proteins and mammalian SWI/SNF complex"

Materials and Methods

**Protein expression, purification, and labeling:** A list of proteins used in the study is provided in Table S4 along with their amino acid sequences. Codon-optimized proteins used in this work were gene-synthesized by GenScript USA Inc. (Piscataway, NJ, USA) and cloned into pET His6 MBP N10 TEV LIC cloning vector (2C-T) [was a gift from Scott Gradia (Addgene plasmid # 29706)]. Proteins were expressed, purified, and fluorescently labeled as described in our earlier work [1]. FOXG1$^{N-IDR}$ was expressed in BL21-CodonPlus (DE3)-RIPL competent cells and was purified using the same protocol as other constructs. All recombinant proteins contained three exogenous amino acids (SNI) at their N-termini after TEV cleavage.

**Cell culture:** The HEK293T cells were cultured at 37°C and 5% $CO_2$ in Dulbecco's Modified Eagle's Medium (Gibco™ 11965092) supplemented with 10% fetal bovine serum (Gibco™ A3160501). To initiate transfection, 20,000 cells were seeded in Nunc™ Lab-Tek™ II chambered coverglass (8 wells). After 24 hours, Lipofectamine 2000 reagent (Thermofisher 11668030) was used to transfect the cells with 0.5 µg plasmid, according to the manufacturer's protocol. The cells were imaged after 20-24 hours of transfection. Colocalization experiments were carried out with 0.5 µg plasmid for each construct. Table S5 provides a complete list of plasmids used for protein expression in HEK293T cells.

**Fluorescence imaging:** To facilitate live-cell imaging, the cells were moved to FluoroBrite DMEM Media (Thermofisher A1896701) containing 10% FBS and Hoechst33342 dye (Thermofisher H3570; 1µg/ml) one hour before the imaging. Imaging was carried out using either a Zeiss LSM710 laser scanning confocal microscope (Plan-Apochromat 63x/1.4 oil DIC M27) or Q2 laser scanning confocal microscope (ISS Inc., 63X objective), with the cells being maintained at 37 °C on a temperature and $CO_2$-controlled stage. In experiments involving the optoFUS$^{PLD-NLS}$ constructs, the Cry2 homo-oligomerization domain (previously characterized by Shin et al., 2017) was fused to FUS$^{PLD-NLS}$ to achieve light-dependent protein condensation in live cells. OptoFUS$^{PLD-NLS}$ droplets were formed by exposing the cells to blue light (488 nm) for a minute during imaging. Image processing was carried out using FIJI and CellProfiler [3,4].

**In vitro phase separation experiments:** The proteins were buffer exchanged into a 25 mM Tris-HCl buffer (pH 7.5) containing 125 mM NaCl at room temperature. After preparing the samples at the desired protein concentrations, TEV protease (TEV: protein = 1:25 v/v) was added and the mixture was incubated for 1 hour at 30 °C to cleave the $His_6$-MBP-N10 tag. 10% Ficoll PM70 was used as a crowder for some samples as mentioned in the appropriate data figures. Next, 4 µl of the sample was placed in the center of a microscope glass slide that was fitted with a custom-made containment, created using the broad end of a plastic pipette tip, and sealed onto the slide. The top of the chamber was then sealed with parafilm to prevent evaporation, and the samples were incubated for 45-60 minutes at room temperature. Finally, the samples were imaged using a Zeiss Primovert inverted iLED microscope (40x objective) or a Zeiss LSM710 laser scanning confocal microscope (Plan-Apochromat 63x/1.4 oil DIC M27). Microscopy images were recorded and processed using ZEN (blue, v2.3) for Zeiss Primovert and ZEN (SP5 2012 Black) for Zeiss LSM710.

**Partition coefficient analysis:** For partition coefficient analysis, a Zeiss LSM710 laser scanning confocal microscope (Plan-Apochromat 63x/1.4 oil DIC M27) was used to record condensate images. Phase-separated condensates were prepared following the procedure outlined above with a trace amount (~ 1-2%) of fluorescently labeled proteins. Confocal images were collected 1 hour after sample preparation. Using CellProfiler, droplets were segmented and the mean intensity within each droplet ($I_{dense}$) was determined. Additionally, for each image, five spots outside the condensates were randomly selected to estimate the background signal, and the mean intensity of the external dilute phase was calculated ($I_{dilute}$). Finally, the partition coefficient ($k$) was calculated by taking the ratio of the two intensities ($k = I_{dense}/I_{dilute}$). For determining the enrichment coefficients in cells, images were imported to CellProfiler, and GFP-PLD condensates were segmented. After segmentation, the mean intensity of the GFP signal and the mean intensity of the mCherry signal within the condensates were obtained. 5-8 spots within the nucleus surrounding the GFP-PLD condensates were randomly selected to estimate the background signal. Enrichment coefficient analysis was then performed as above. For in vitro experiments, a total of 100-300 condensates were analyzed, whereas, for cellular experiments, a total of 30-80 condensates were analyzed for each PLD construct.

**Estimation of saturation concentration for PLDs in cells:** Images of cells expressing GFP-tagged prion-like domains (PLDs) or variants were captured using a laser scanning confocal microscope (Q2-ISS Inc., 63X objective) with fixed imaging parameters across samples. Images were imported into CellProfiler, and nuclei of the cells were segmented using Hoechst intensity. After segmentation, mean GFP intensity was obtained within the segmented region as a proxy for nuclear protein concentration and converted to absolute concentration using the GFP standard curve.

**Fluorescent recovery after photobleaching (FRAP) analysis:** For FRAP experiments, a circular region of interest was bleached with 100% power for ~1-2 seconds which was followed by an imaging scan for 60 seconds. The recorded AlexaFluor488-labeled probe intensity or GFP intensity values from the bleached ROI were then corrected for photofading by normalizing them to an unbleached reference condensate. Multiple FRAP recovery curves were averaged for each sample and plotted as a function of time using OriginPro (2018b). The dimension of the bleaching ROI was constant across samples. For each time point of the FRAP recovery curve, the standard deviation of the intensity values from the three or more FRAP curves was taken as the uncertainty.

**Bead Halo Assay (BHA):** 20 µl of HisPur™ Ni-NTA Magnetic Beads (Thermo Fisher Scientific 88831) slurry was resuspended in 480 µl of buffer (25 mM Tris-HCl, pH 7.5, 125 mM NaCl), mixed by inversion of the tube, and then centrifuged at 125 x g. The supernatant was removed, and the wash step was repeated two additional times. The beads were finally resuspended in 200 µl buffer, which was subsequently used as a working stock solution. For the experimental setup, 1 µl of beads was diluted in 3 µl of buffer and 0.5 µl of 2.5 µM AlexaFluor488-labeled scaffold. These proteins contained a hexahistidine ($His_6$) tag, which enabled their immobilization on the bead surface via Ni-$His_6$ interactions. The scaffolds also contained an MBP solubility tag to prevent their surface condensation [5]. The scaffold-coated beads were allowed to equilibrate for 15 minutes, following which 0.5 µl of 2.5 µM of AlexaFluor594-labeled client ($FUS^{PLD}$) was added to achieve a final concentration of 250 nM of both the scaffold PLD and the client PLD. The samples were incubated for 15 minutes and imaged using a laser scanning confocal microscope (Q2 laser scanning microscope, 63X objective). For estimating the relative enrichment of $FUS^{PLD}$ on the scaffold-coated beads, a ratio of the mean fluorescence intensity of the client (AlexaFluor594) on the bead to the mean fluorescence intensity of the scaffold (AlexaFluor488) was taken and normalized to the labeling efficiency. This normalized data was subsequently defined as the client enrichment score and compared across samples (**Fig. 6c**). For our attempt to estimate a binding curve for heterotypic PLD complexation, the client ($FUS^{PLD}$) concentration was continuously varied as indicated in **Fig. S16**, keeping the scaffold concentration fixed at 250 nM.

### Supplementary Tables

| | ARID1A$^{PLD}$ | ARID1B$^{PLD}$ | BRG1$^{PLD}$ | SS18$^{PLD}$ | SMARCC1$^{PLD}$ | FUS$^{PLD}$ | hPol II CTD$^{1-30}$ | EWSR1$^{PLD}$ | TAF15$^{PLD}$ |
|---|---|---|---|---|---|---|---|---|---|
| Ala (A) | 63 | 60 | 25 | 3 | 12 | 4 | 1 | 27 | 2 |
| Arg (R) | 12 | 11 | 9 | 7 | 2 | 0 | 1 | 1 | 6 |
| Asn (N) | 14 | 8 | 8 | 25 | 4 | 5 | 5 | 4 | 10 |
| Asp (D) | 7 | 5 | 8 | 9 | 1 | 2 | 0 | 5 | 14 |
| Cys (C) | 1 | 3 | 0 | 0 | 0 | 1 | 0 | 0 | 0 |
| Gln (Q) | 88 | 49 | 27 | 72 | 18 | 37 | 1 | 47 | 35 |
| Glu (E) | 3 | 2 | 4 | 5 | 1 | 0 | 1 | 1 | 6 |
| Gly (G) | 83 | 92 | 56 | 58 | 20 | 35 | 2 | 25 | 28 |
| His (H) | 13 | 8 | 11 | 11 | 10 | 0 | 0 | 0 | 5 |
| Ile (I) | 1 | 2 | 3 | 1 | 4 | 0 | 0 | 0 | 0 |
| Leu (L) | 15 | 10 | 12 | 3 | 5 | 0 | 0 | 1 | 0 |
| Lys (K) | 4 | 3 | 7 | 0 | 0 | 0 | 0 | 1 | 1 |
| Met (M) | 10 | 22 | 26 | 31 | 12 | 1 | 1 | 2 | 2 |
| Phe (F) | 6 | 3 | 1 | 0 | 0 | 0 | 0 | 0 | 0 |
| Pro (P) | 87 | 51 | 90 | 63 | 48 | 11 | 60 | 30 | 4 |
| Ser (S) | 66 | 42 | 31 | 24 | 5 | 44 | 79 | 40 | 36 |
| Thr (T) | 18 | 6 | 10 | 4 | 3 | 9 | 31 | 39 | 4 |
| Trp (W) | 1 | 1 | 1 | 0 | 0 | 0 | 0 | 0 | 0 |
| Tyr (Y) | 29 | 22 | 5 | 32 | 2 | 24 | 30 | 37 | 26 |
| Val (V) | 3 | 4 | 6 | 4 | 3 | 0 | 0 | 5 | 1 |

**Table S1:** Amino acid composition (in percentage) for each of the prion-like domains (PLDs) used in this study. The red color intensity scales linearly with the percentage of a given amino acid.

| | ARID1A$^{PLD}$ | ARID1B$^{PLD}$ | BRG1$^{PLD}$ | SS18$^{PLD}$ | SMARCC1$^{PLD}$ | FUS$^{PLD}$ | Pol II CTD$^{1-30}$ | EWSR1$^{PLD}$ | TAF15$^{PLD}$ |
|---|---|---|---|---|---|---|---|---|---|
| Ala (A) | 63 | 60 | 25 | 3 | 12 | 4 | 1 | 27 | 2 |
| Arg (R) | 12 | 11 | 9 | 7 | 2 | 0 | 1 | 1 | 6 |
| Asn (N) | 14 | 8 | 8 | 25 | 4 | 5 | 5 | 4 | 10 |
| Asp (D) | 7 | 5 | 8 | 9 | 1 | 2 | 0 | 5 | 14 |
| Cys (C) | 1 | 3 | 0 | 0 | 0 | 1 | 0 | 0 | 0 |
| Gln (Q) | 88 | 49 | 27 | 72 | 18 | 37 | 1 | 47 | 35 |
| Glu (E) | 3 | 2 | 4 | 5 | 1 | 0 | 1 | 1 | 6 |
| Gly (G) | 83 | 92 | 56 | 58 | 20 | 35 | 2 | 25 | 28 |
| His (H) | 13 | 8 | 11 | 11 | 10 | 0 | 0 | 0 | 5 |
| Ile (I) | 1 | 2 | 3 | 1 | 4 | 0 | 0 | 0 | 0 |
| Leu (L) | 15 | 10 | 12 | 3 | 5 | 0 | 0 | 1 | 0 |
| Lys (K) | 4 | 3 | 7 | 0 | 0 | 0 | 0 | 1 | 1 |
| Met (M) | 10 | 22 | 26 | 31 | 12 | 1 | 1 | 2 | 2 |
| Phe (F) | 6 | 3 | 1 | 0 | 0 | 0 | 0 | 0 | 0 |
| Pro (P) | 87 | 51 | 90 | 63 | 48 | 11 | 60 | 30 | 4 |
| Ser (S) | 66 | 42 | 31 | 24 | 5 | 44 | 79 | 40 | 36 |
| Thr (T) | 18 | 6 | 10 | 4 | 3 | 9 | 31 | 39 | 4 |
| Trp (W) | 1 | 1 | 1 | 0 | 0 | 0 | 0 | 0 | 0 |
| Tyr (Y) | 29 | 22 | 5 | 32 | 2 | 24 | 30 | 37 | 26 |
| Val (V) | 3 | 4 | 6 | 4 | 3 | 0 | 0 | 5 | 1 |

**Table S2:** Number of amino acids in each of the prion-like domains (PLDs) used in this study. The red color intensity scales linearly with the number of a given amino acid.

| | ARID1A$^{PLD}$ | ARID1B$^{PLD}$ | BRG1$^{PLD}$ | SS18$^{PLD}$ | SMARCC1$^{PLD}$ | FUS$^{PLD}$ | Pol II CTD[30] | EWSR1$^{PLD}$ | TAF15$^{PLD}$ |
|---|---|---|---|---|---|---|---|---|---|
| Aromatic stickers (FYW) | 36 | 26 | 7 | 32 | 2 | 24 | 30 | 37 | 26 |
| Positive Charges (RK) | 16 | 14 | 16 | 7 | 2 | 0 | 1 | 2 | 7 |
| Negative charges (DE) | 10 | 7 | 12 | 14 | 2 | 2 | 1 | 6 | 20 |
| Hydrophobi c (ILMV) | 29 | 38 | 47 | 39 | 24 | 1 | 1 | 8 | 3 |
| Arginines +Aromatic (R+FYW) | 48 | 37 | 16 | 39 | 4 | 24 | 31 | 38 | 32 |
| NCPR | 0.011 | 0.017 | 0.012 | -0.020 | 0.000 | -0.011 | 0.000 | -0.015 | -0.012 |
| Length | 524 | 404 | 340 | 352 | 150 | 173 | 212 | 265 | 180 |
| Sticker density ((R+FYW)/ Length) | 0.09 | 0.09 | 0.05 | 0.11 | 0.03 | 0.14 | 0.15 | 0.14 | 0.18 |

**Table S3:** Amino acid features of the prion-like domains employed in this work. Net charge per residue (NCPR) values were obtained using the CIDER tool [6]. The red color intensity scales linearly with the value of a given parameter.

| Protein | Purpose | Sequence |
|---|---|---|
| BRG1$^{PLD}$ | Purification/ live-cell imaging | MSTPDPPLGGTPRPGPSPGPGPSPGAMLGPSPGPSPGSAHSMMGPS<br>PGPPSAGHPIPTQGPGGYPQDNMHQMHKPMESMHEKGMSDDPRYNQ<br>MKGMGMRSGGHAGMGPPPSPMDQHSQGYPSPLGGSEHASSPVPAS<br>GPSSGPQMSSGPGGAPLDGADPQALGQQNRGPTPFNQNQLHQLRAQI<br>MAYKMLARGQPLPDHLQMAVQGKRPMPGMQQQMPTLPPPSVSATGP<br>GPGPGPGPGPGPGPAPPNYSRPHGMGGPNMPPPGPSGVPPGMPGQ<br>PPGGPPKPWPEGPMANAAAPTSTPQKLIPPQPTGRPSPAPPAVPPAAS<br>PVMPPQTQSPGQPAQPA |
| BRG1$^{PLD}$ 2C | Purification/ Cys-maleimide conjugation for site-specific protein labeling | MCSTPDPPLGGTPRPGPSPGPGPSPGAMLGPSPGPSPGSAHSMMGP<br>SPGPPSAGHPIPTQGPGGYPQDNMHQMHKPMESMHEKGMSDDPRYN<br>QMKGMGMRSGGHAGMGPPPSPMDQHSQGYPSPLGGSEHASSPVPA<br>SGPSSGPQMSSGPGGAPLDGADPQALGQQNRGPTPFNQNQLHQLRA<br>QIMAYKMLARGQPLPDHLQMAVQGKRPMPGMQQQMPTLPPPSVSATG<br>PGPGPGPGPGPGPGPAPPNYSRPHGMGGPNMPPPGPSGVPPGMPG<br>QPPGGPPKPWPEGPMANAAAPTSTPQKLIPPQPTGRPSPAPPAVPPAA<br>SPVMPPQTQSPGQPAQPA |
| ARID1A$^{PLD}$ | Purification/ Cys-maleimide conjugation for site-specific protein labeling / live-cell imaging | MSNGGGGGGGAGSGGGPGAEPDLKNSNGNAGPRPALNNNLTEPPGG<br>GGGGSSDGVGAPPHSAAAALPPPAYGFGQPYGRSPSAVAAAAAAVFH<br>QQHGGQQSPGLAALQSGGGGGLEPYAGPQQNSHDHGFPNHQYNSYY<br>PNRSAYPPPAPAYALSSPRGGTPGSGAAAAAGSKPPPSSSASASSSSS<br>SFAQQRFGAMGGGGPSAAGGGTPQPTATPTLNQLLTSPSSARGYQGY<br>PGGDYSGGPQDGGAGKGPADMASQCWGAAAAAAAAAAASGGAQQR<br>SHHAPMSPGSSGGGGQPLARTPQPSSPMDQMGKMRPQPYGGTNPYS<br>QQQGPPSGPQQGHGYPGQPYGSQTPQRYPMTMQGRAQSAMGGLSY<br>TQQIPPYGQQGPSGYGQQGQTPYYNQQSPHPQQQQPPYSQQPPSQT<br>PHAQPSYQQQPQSQPPQLQSSQPPYSQQPSQPPHQQSPAPYPSQQS<br>TTQQHPQSQPPYSQPQAQSPYQQQQPQQPAPSTLSQQAAYPQPQSQ<br>QSQQTAYSQQRFPPPQ |

| | | |
|---|---|---|
| ARID1B$^{PLD}$ | Purification/ Cys-maleimide conjugation for site-specific protein labeling / live-cell imaging | MNNYYGSAAPASGGPGGRAGPCFDQHGGQQSPGMGMMHSASAAAA<br>GAPGSMDPLQNSHEGYPNSQCNHYPGYSRPGAGGGGGGGGGGGG<br>SGGGGGGGGAGAGGAGAGAVAAAAAAAAAAAAGGGGGGGYGGSSAG<br>YGVLSSPRQQGGGMMMGPGGGGAASLSKAAAGSAAGGFQRFAGQN<br>QHPSGATPTLNQLLTSPSPMMRSYGGSYPEYSSPSAPPPPPSQPQSQA<br>AAAGAAAGGQQAAAGMGLGKDMGAQYAAASPAWAAAQQRSHPAMSP<br>GTPGPTMGRSQGSPMDPMVMKRPQLYGMGSNPHSQPQQSSPYPGG<br>SYGPPGPQRYPIGIQGRTPGAMAGMQYPQQQMPPQYGQQGVSGYCQ<br>QGQQPYYSQQPQPPHLPPQAQYLPSQSQQRYQPQQDMSQ |
| SS18$^{PLD}$ | Purification/ live-cell imaging | MNQNMQSLLPAPPTQNMPMGPGGMNQSGPPPPPRSHNMPSDGMVG<br>GGPPAPHMQNQMNGQMPGPNHMPMQGPGPNQLNMTNSSMNMPSSS<br>HGSMGGYNHSVPSSQSMPVQNQMTMSQGQPMGNYGPRPNMSMQPN<br>QGPMMHQQPPSQQYNMPQGGGQHYQGQQPPMGMMGQVNQGNHM<br>MGQRQIPPYRPPQQGPPQQYSGQEDYYGDQYSHGGQGPPEGMNQQ<br>YYPDGHNDYGYQQPSYPEQGYDRPYEDSSQHYYEGGNSQYGQQQDA<br>YQGPPPQQGYPPQQQQYPGQQGYPGQQQGYGPSQGGPGPQYPNYP<br>QGQGQQYGGYRPTQPGPPQPPQQRPYGYDQGQYGNYQQ |
| SS18$^{PLD}$ 2C | Purification/ Cys-maleimide conjugation for site-specific protein labeling | MCNQNMQSLLPAPPTQNMPMGPGGMNQSGPPPPPRSHNMPSDGMV<br>GGGPPAPHMQNQMNGQMPGPNHMPMQGPGPNQLNMTNSSMNMPS<br>SSHGSMGGYNHSVPSSQSMPVQNQMTMSQGQPMGNYGPRPNMSMQ<br>PNQGPMMHQQPPSQQYNMPQGGGQHYQGQQPPMGMMGQVNQGNH<br>MMGQRQIPPYRPPQQGPPQQYSGQEDYYGDQYSHGGQGPPEGMNQ<br>QYYPDGHNDYGYQQPSYPEQGYDRPYEDSSQHYYEGGNSQYGQQQD<br>AYQGPPPQQGYPPQQQQYPGQQGYPGQQQGYGPSQGGPGPQYPNY<br>PQGQGQQYGGYRPTQPGPPQPPQQRPYGYDQGQYGNYQQ |
| SMARCC1$^{PLD}$ | Live cell imaging | QQMEQQQHGQNPQQAHQHSGGPGLAPLGAAGHPGMMPHQQPPPYP<br>LMHHQMPPPHPPQPGQIPGPGSMMPGQHMPGRMIPTVAANIHPSGSG<br>PTPPGMPPMPGNILGPRVPLTAPNGMYPPPPQQQPPPPPPADGVPPPP<br>APGPPASAAP |
| FUS$^{PLD}$ | Purification/ live-cell imaging/ optogenetic constructs | MASNDYTQQATQSYGAYPTQPGQGYSQQSSQPYGQQSYSGYSQSTD<br>TSGYGQSSYSSYGQSQNSYGTQSTPQGYGSTGGYGSSQSSQSSYGQ<br>QSSYPGYGQQPAPSSTSGSYGSSSQSSSYGQPQSGSYSQQPSYGGQ<br>QQSYGQQQSYNPPQGYGQQNQYNSSSGGGGGGGGG |
| FUS$^{PLD}$ S86C | Purification/ Cys-maleimide conjugation for site-specific protein labeling | MASNDYTQQATQSYGAYPTQPGQGYSQQSSQPYGQQSYSGYSQSTD<br>TSGYGQSSYSSYGQSQNSYGTQSTPQGYGSTGGYGSSQSCQSSYGQ |

| | | |
|---|---|---|
| | | QSSYPGYGQQPAPSSTSGSYGSSSQSSSYGQPQSGSYSQQPSYGGQ<br>QQSYGQQQSYNPPQGYGQQNQYNSSSGGGGGGGG |
| EWSR1$^{\text{PLD}}$ | Live cell imaging | MASTDYSTYSQAAAQQGYSAYTAQPTQGYAQTTQAYGQQSYGTYGQP<br>TDVSYTQAQTTATYGQTAYATSYGQPPTGYTTPTAPQAYSQPVQGYGT<br>GAYDTTTATVTTTQASYAAQSAYGTQPAYPAYGQQPAATAPTRPQDGN<br>KPTETSQPQSSTGGYNQPSLGYGQSNYSYPQVPGSYPMQPVTAPPSY<br>PPTSYSSTQPTSYDQSSYSQQNTYGQPSSYGQQSSYGQQSSYGQQPP<br>TSYPPQTGSYSQAPSQYSQQSSSYGQQS |
| TAF15$^{\text{PLD}}$ | Live cell imaging | SDSGSYSQSGGEQQSYSSYGNQGSQGYGQTPQGYSGYGQTTDSSYG<br>QNYGGYSGYGQNQSGYSQSYGSYENQKQSSYGQQSYNNQGQQNTE<br>SSGGQGGRAPSYGQSDYGQQDSYDQQSGYDQHQGSYDEQSNYQQH<br>DSYNQNQQSYHSQRENYSHHTQDDRRDVSRYGEDNRGYGGSQGGG<br>RGRGGYDKDGRGPMTGSSGGDRG |
| RNA Pol II CTD$^{\text{1-30}}$ | Live cell imaging | YSPTSPAYEPRSPGGYTPQSPSYSPTSPSYSPTSPSYSPTSPNYSPTSP<br>SYSPTSPSYSPTSPSYSPTSPSYSPTSPSYSPTSPSYSPTSPSYSPTSP<br>SYSPTSPSYSPTSPSYSPTSPSYSPTSPSYSPTSPSYSPTSPSYSPTSP<br>SYSPTSPNYSPTSPNYTPTSPSYSPTSPSYSPTSPNYTPTSPNYSPTSP<br>SYSPTSPSYSPTSPS |
| RNA Pol II CTD$^{\text{1-30}}$ 2C | Purification/ Cys-maleimide conjugation for site-specific protein labeling | MCYSPTSPAYEPRSPGGYTPQSPSYSPTSPSYSPTSPSYSPTSPNYSP<br>TSPSYSPTSPSYSPTSPSYSPTSPSYSPTSPSYSPTSPSYSPTSPSYSPT<br>SPSYSPTSPSYSPTSPSYSPTSPSYSPTSPSYSPTSPSYSPTSPSYSPT<br>SPSYSPTSPNYSPTSPNYTPTSPSYSPTSPSYSPTSPNYTPTSPNYSPT<br>SPSYSPTSPSYSPTSPS |
| FOXG1$^{\text{N-IDR}}$ | Live cell imaging | MLDMGDRKEVKMIPKSSFSINSLVPEAVQN DNHHASHGHHNSHHPQH<br>HHHHHHHHHHPPPPAPQPPPPPPQQQPPPPPPPAPQPPQTRGAPAAD<br>DDKGPQQLLLPPPPPPPPAAALDGAKADGLGGKGEPGGGPGELAPVG<br>PDEKEKGAGAGGEEKKGAGEGGKDGEGGKEGEKKNGKYE |
| FOXG1$^{\text{N-IDR}}$ 2C | Purification/ Cys-maleimide conjugation for site-specific protein labeling / | MCLDMGDRKEVKMIPKSSFSINSLVPEAVQN DNHHASHGHHNSHHPQ<br>HHHHHHHHHHHPPPPAPQPPPPPPQQQPPPPPPPAPQPPQTRGAPAA<br>DDDKGPQQLLLPPPPPPPPAAALDGAKADGLGGKGEPGGGPGELAPV<br>GPDEKEKGAGAGGEEKKGAGEGGKDGEGGKEGEKKNGKYE |
| His-MBP-TEV-cys | Purification/ Cys-maleimide conjugation for site-specific protein labeling | GGGCGGG |

| | | |
|---|---|---|
| BRG1$^{PLD\ Aro+}$ | Live cell imaging | MSTPDPPLGGTYRPGPSPGPGPSPGAMLGPSPGPSYGSAHSMMGPS<br>PGPYSAGHPIPTQGPGGYYQDNMHQMHKPMESMHEKGMSDDPRYNQ<br>MKGMGMRSGGHAGMGPYPSPMDQHSQGYPSPLGGSEHASSPVPAS<br>GPSSGPQMSSGPGGAYLDGADPQALGQQNRGPTPFNQNQLHQLRAQI<br>MAYKMLARGQYLPDHLQMAVQGKRPMPGMQQQMYTLPPPSVSATGY<br>GPGPGPGPGPGYGPAPPNYSRYHGMGGPNMPPYGPSGVPPGMPGQ<br>PPGGPPKPWYEGPMANAAAPTSTPQKLIYPQPTGRPSYAPPAVPYAAS<br>PVMPPQTQSYGQPAQPA |
| BRG1$^{PLD\ Aro++}$ | Live cell imaging | MSTPDYPLGGTYRPGYSPGYGPSYGAMLGPSPGPSYGSAHSMMGPS<br>YGPYSAGHPIYTQGPGGYYQDNMHQMHKPMESMHEKGMSDDYRYNQ<br>MKGMGMRSGGHAGMGPYPSYMDQHSQGYPSYLGGSEHASSPVPAS<br>GPSSGYQMSSGPGGAYLDGADPQALGQQNRGYTPFNQNQLHQLRAQI<br>MAYKMLARGQYLPDHLQMAVQGKRYMPGMQQQMYTLPYPSVSATGY<br>GPGPGPGYGPGYGPAYPNYSRYHGMGGPNMYPYGPSGVYPGMYGQ<br>PYGGPPKPWYEGPMANAAAYTSTPQKLIYPQYTGRPSYAPYAVPYAAS<br>PVMYPQTQSYGQPAQYA |
| FUS$^{2XPLD}$ | Live cell imaging | MASNDYTQQATQSYGAYPTQPGQGYSQQSSQPYGQQSYSGYSQSTD<br>TSGYGQSSYSSYGQSQNSYGTQSTPQGYGSTGGYGSSQSSQSSYGQ<br>QSSYPGYGQQPAPSSTSGSYGSSSQSSSYGQPQSGSYSQQPSYGGQ<br>QQSYGQQQSYNPPQGYGQQNQYNSSSGGGGGGGGGMASNDYTQQA<br>TQSYGAYPTQPGQGYSQQSSQPYGQQSYSGYSQSTDTSGYGQSSYS<br>SYGQSQNSYGTQSTPQGYGSTGGYGSSQSSQSSYGQQSSYPGYGQQ<br>PAPSSTSGSYGSSSQSSSYGQPQSGSYSQQPSYGGQQQSYGQQQSY<br>NPPQGYGQQNQYNSSSGGGGGGGGG |
| FUS$^{2XPLD\ halfYtoS}$ | Live cell imaging | MASNDYTQQATQSYGAYPTQPGQGYSQQSSQPYGQQSYSGYSQSTD<br>TSGYGQSSYSSYGQSQNSYGTQSTPQGYGSTGGYGSSQSSQSSYGQ<br>QSSYPGYGQQPAPSSTSGSYGSSSQSSSYGQPQSGSYSQQPSYGGQ<br>QQSYGQQQSYNPPQGYGQQNQYNSSSGGGGGGGGGMASNDSTQQA<br>TQSSGASPTQPGQGSSQQSSQPSGQQSSGSSQSTDTSGSGQSSSS<br>SSGQSQNSSGTQSTPQGSGSTGGSGSSQSSQSSSGQQSSSPGSGQQ<br>PAPSSTSGSSGSSSQSSSSGQPQSGSSSQQPSSGGQQQSSGQQQSS<br>NPPQGSGQQNQSNSSSGGGGGGGGG |
| BRG1$^{Folded}$ | Live cell imaging | EYGVSQALARGLQSYYAVAHAVTERVDKQSALMVNGVLKQYQIKGLEW<br>LVSLYNNNLNGILADEMGLGKTIQTIALITYLMEHKRINGPFLIIVPLSTLSN |

| | | |
|---|---|---|
| | | WAYEFDKWAPSVVKVSYKGSPAARRAFVPQLRSGKFNVLLTTYEYIIKD KHILAKIRWKYMIVDEGHRMKNHHCKLTQVLNTHYVAPRRLLLTGTPLQ NKLPELWALLNFLLPTIFKSCSTFEQWFNAPFAMTGEKVDLNEEETILIIR RLHKVLRPFLLRRLKKEVEAQLPEKVEYVIKCDMSALQRVLYRHMQAKG VLLTDGSEKDKKGKGGTKTLMNTIMQLRKICNHPYMFQHIEESFSEHLG FTGGIVQGLDLYRASGKFELLDRILPKLRATNHKVLLFCQMTSLMTIMED YFAYRGFKYLRLDGTTKAEDRGMLLKTFNEPGSEYFIFLLSTRAGGLGL NLQSADTVIIFDSDWNPHQDLQAQDRAHRIGQQNEVRVLRLCTVNSVEE KILAAAKYKLNVDQKVIQAGM |
| BRG1 | Live-cell imaging | MSTPDPPLGGTPRPGPSPGPGPSPGAMLGPSPGPSPGSAHSMMGPS PGPPSAGHPIPTQGPGGYPQDNMHQMHKPMESMHEKGMSDDPRYNQ MKGMGMRSGGHAGMGPPPSPMDQHSQGYPSPLGGSEHASSPVPAS GPSSGPQMSSGPGGAPLDGADPQALGQQNRGPTPFNQNQLHQLRAQI MAYKMLARGQPLPDHLQMAVQGKRPMPGMQQQMPTLPPPSVSATGP GPGPGPGPGPGPGPAPPNYSRPHGMGGPNMPPPGPSGVPPGMPGQ PPGGPPKPWPEGPMANAAAPTSTPQKLIPPQPTGRPSPAPPAVPPAAS PVMPPQTQSPGQPAQPAPMVPLHQKQSRITPIQKPRGLDPVEILQEREY RLQARIAHRIQELENLPGSLAGDLRTKATIELKALRLLNFQRQLRQEVVV CMRRDTALETALNAKAYKRSKRQSLREARITEKLEKQQKIEQERKRRQK HQEYLNSILQHAKDFKEYHRSVTGKIQKLTKAVATYHANTEREQKKENE RIEKERMRRLMAEDEEGYRKLIDQKKDKRLAYLLQQTDEYVANLTELVR QHKAAQVAKEKKKKKKKKKAENAEGQTPAIGPDGEPLDETSQMSDLPV KVIHVESGKILTGTDAPKAGQLEAWLEMNPGYEVAPRSDSEESGSEEEE EEEEEEQPQAAQPPTLPVEEKKKIPDPDSDDVSEVDARHIIENAKQDVD DEYGVSQALARGLQSYYAVAHAVTERVDKQSALMVNGVLKQYQIKGLE WLVSLYNNNLNGILADEMGLGKTIQTIALITYLMEHKRINGPFLIIVPLSTLS NWAYEFDKWAPSVVKVSYKGSPAARRAFVPQLRSGKFNVLLTTYEYIIK DKHILAKIRWKYMIVDEGHRMKNHHCKLTQVLNTHYVAPRRLLLTGTPL QNKLPELWALLNFLLPTIFKSCSTFEQWFNAPFAMTGEKVDLNEEETILII RRLHKVLRPFLLRRLKKEVEAQLPEKVEYVIKCDMSALQRVLYRHMQAK GVLLTDGSEKDKKGKGGTKTLMNTIMQLRKICNHPYMFQHIEESFSEHL GFTGGIVQGLDLYRASGKFELLDRILPKLRATNHKVLLFCQMTSLMTIME DYFAYRGFKYLRLDGTTKAEDRGMLLKTFNEPGSEYFIFLLSTRAGGLG LNLQSADTVIIFDSDWNPHQDLQAQDRAHRIGQQNEVRVLRLCTVNSVE EKILAAAKYKLNVDQKVIQAGMFDQKSSSHERRAFLQAILEHEEQDESR HCSTGSGSASFAHTAPPPAGVNPDLEEPPLKEEDEVPDDETVNQMIAR |

| | | |
|---|---|---|
| | | HEEEFDLFMRMDLDRRREEARNPKRKPRLMEEDELPSWIIKDDAEVERL<br>TCEEEEEEKMFGRGSRHRKEVDYSDSLTEKQWLKAIEEGTLEEIEEEVRQ<br>KKSSRKRKRDSDAGSSTPTTSTRSRDKDDESKKQKKRGRPPAEKLSPN<br>PPNLTKKMKKIVDAVIKYKDSSSGRQLSEVFIQLPSRKELPEYYELIRKPV<br>DFKKIKERIRNHKYRSLNDLEKDVMLLCQNAQTFNLEGSLIYEDSIVLQSV<br>FTSVRQKIEKEDDSEGEESEEEEEGEEEGSESESRSVKVKIKLGRKEKA<br>QDRLKGGRRRPSRGSRAKPVVSDDDSEEEQEEDRSGSGSEED |
| ARID1A$^{PLD\ 30QtoG}$ | Live cell imaging | MSNGGGGGGGAGSGGGPGAEPDLKNSNGNAGPRPALNNNLTEPPGG<br>GGGGSSDGVGAPPHSAAAALPPPAYGFGQPYGRSPSAVAAAAAAVFH<br>QQHGGQQSPGLAALQSGGGGGLEPYAGPQQNSHDHGFPNHQYNSYY<br>PNRSAYPPPAPAYALSSPRGGTPGSGAAAAAGSKPPPSSSASASSSSS<br>SFAQQRFGAMGGGGPSAAGGGTPQPTATPTLNQLLTSPSSARGYQGY<br>PGGDYSGGPQDGGAGKGPADMASQCWGAAAAAAAAAAASGGAQQR<br>SHHAPMSPGSSGGGGQPLARTPQPSSPMDQMGKMRPQPYGGTNPYS<br>GGGGPPSGPQQGHGYPGQPYGSQTPQRYPMTMQGRAQSAMGGLSY<br>TQQIPPYGQQGPSGYGQQGQTPYYNQQSPHPGGGGPPYSQQPPSQT<br>PHAQPSYGGGPGSGPPGLGSSQPPYSQQPSQPPHQQSPAPYPSGGS<br>TTGGHPQSQPPYSQPQAQSPYGGGGPGGPAPSTLSQQAAYPGPGSG<br>GSGGTAYSQQRFPPPQ |
| ARID1A$^{PLD\ YtoS}$ | Live cell imaging | MSNGGGGGGGAGSGGGPGAEPDLKNSNGNAGPRPALNNNLTEPPGG<br>GGGGSSDGVGAPPHSAAAALPPPASGFGQPSGRSPSAVAAAAAAVFH<br>QQHGGQQSPGLAALQSGGGGGLEPSAGPQQNSHDHGFPNHQSNSSS<br>PNRSASPPPAPASALSSPRGGTPGSGAAAAAGSKPPPSSSASASSSSS<br>SFAQQRFGAMGGGGPSAAGGGTPQPTATPTLNQLLTSPSSARGSQGS<br>PGGDSSGGPQDGGAGKGPADMASQCWGAAAAAAAAAAASGGAQQR<br>SHHAPMSPGSSGGGGQPLARTPQPSSPMDQMGKMRPQPSGGTNPSS<br>QQQGPPSGPQQGHGSPGQPSGSQTPQRSPMTMQGRAQSAMGGLSS<br>TQQIPPSGQQGPSGSGQQGQTPSSNQQSPHPQQQQPPSSQQPPSQT<br>PHAQPSSQQQPQSQPPQLQSSQPPSSQQPSQPPHQQSPAPSPSQQS<br>TTQQHPQSQPPSSQPQAQSPSQQQQPQQPAPSTLSQQAASPQPQSQ<br>QSQQTASSQQRFPPPQ |

**Table S4:** List of protein constructs used for recombinant protein expression and purification and live cell imaging in this work. The mutation sites are highlighted in blue.

| Plasmid | Tag | Source plasmid |
|---|---|---|
| BRG1$^{PLD}$ | mEGFP | Synthesized by genscript |
| ARID1A$^{PLD}$ | mEGFP | Synthesized by genscript |
| ARID1B$^{PLD}$ | mEGFP | Synthesized by genscript |
| SS18$^{PLD}$ | mEGFP | Synthesized by genscript |
| FUS$^{PLD}$ | mEGFP | Synthesized by genscript |
| FUS$^{2XPLD}$ | mEGFP | Synthesized by genscript |
| OptoFUS$^{PLD}$ | CRY2-mCherry | Kind gift from Dr. Sreejith Nair |
| BRG1$^{PLD}$ | mCherry | Synthesized by genscript |
| ARID1A$^{PLD}$ | mCherry | Synthesized by genscript |
| ARID1B$^{PLD}$ | mCherry | Synthesized by genscript |
| SS18$^{PLD}$ | mCherry | Synthesized by genscript |
| FUS$^{PLD}$ | mCherry | Synthesized by genscript |
| hRNA Pol II CTD$^{1-30}$ | mCherry | Synthesized by genscript |
| mCherry2-C1 | | Addgene: #54563, a gift from Michael Davidson |
| SMARCC1$^{PLD}$ | GFP | Synthesized by genscript |
| EWSR1$^{PLD}$ | GFP | Synthesized by genscript |
| TAF15$^{PLD}$ | GFP | Synthesized by genscript |
| BRG1$^{PLD\ Aro+}$ | GFP | Synthesized by genscript |
| BRG1$^{PLD\ Aro++}$ | GFP | Synthesized by genscript |
| ARID1A$^{PLD\ YtoS}$ | GFP | Synthesized by genscript |
| ARID1A$^{PLD\ 30QtoG}$ | GFP | Synthesized by genscript |
| FOXG1$^{N-IDR}$ | GFP | Synthesized by genscript |
| FOXG1$^{N-IDR}$ | mCherry | Synthesized by genscript |
| FUS$^{2XPLD\ halfYtoS}$ | GFP | Synthesized by genscript |
| BRG1 | GFP | Addgene # 65391, a gift from Kyle Miller |
| BRG1$^{Folded}$ | GFP | Synthesized by genscript |
| BRG1$^{PLD}$ 2C | His-MBP-N10-Tev | Synthesized by genscript |
| ARID1A$^{PLD}$ | His-MBP-N10-Tev | Synthesized by genscript |
| ARID1B$^{PLD}$ | His-MBP-N10-Tev | Synthesized by genscript |
| SS18$^{PLD}$ 2C | His-MBP-N10-Tev | Synthesized by genscript |
| FUS$^{PLD}$ S86C | His-MBP-N10-Tev | Synthesized by genscript |
| RNA Pol II CTD$^{1-30}$ 2C | His-MBP-N10-Tev | Synthesized by genscript |
| FOXG1$^{N-IDR}$ 2C | His-MBP-N10-Tev | Synthesized by genscript |
| His-MBP-TEV-cys | His-MBP-N10-Tev | Synthesized by genscript |

**Table S5:** List of plasmids used in this work for protein expression in HEK293T cells.

| Side chain chemistry | Amino acid | Color |
|---|---|---|
| Positively charged | Arg, Lys & His | Green |
| Negatively charged | Asp & Glu | Pink |
| Polar uncharged | Ser, Thr, Asn & Gln | Yellow |
| Hydrophobic | Ala, Val, Ile, leu, Met, Phe, Tyr & Trp | Blue |
| Others | Pro, Gly & Cys | Orange |

**Table S6:** Color code for amino acids in Figs.1b, 2a, & 5b.

**Supplementary Figures**

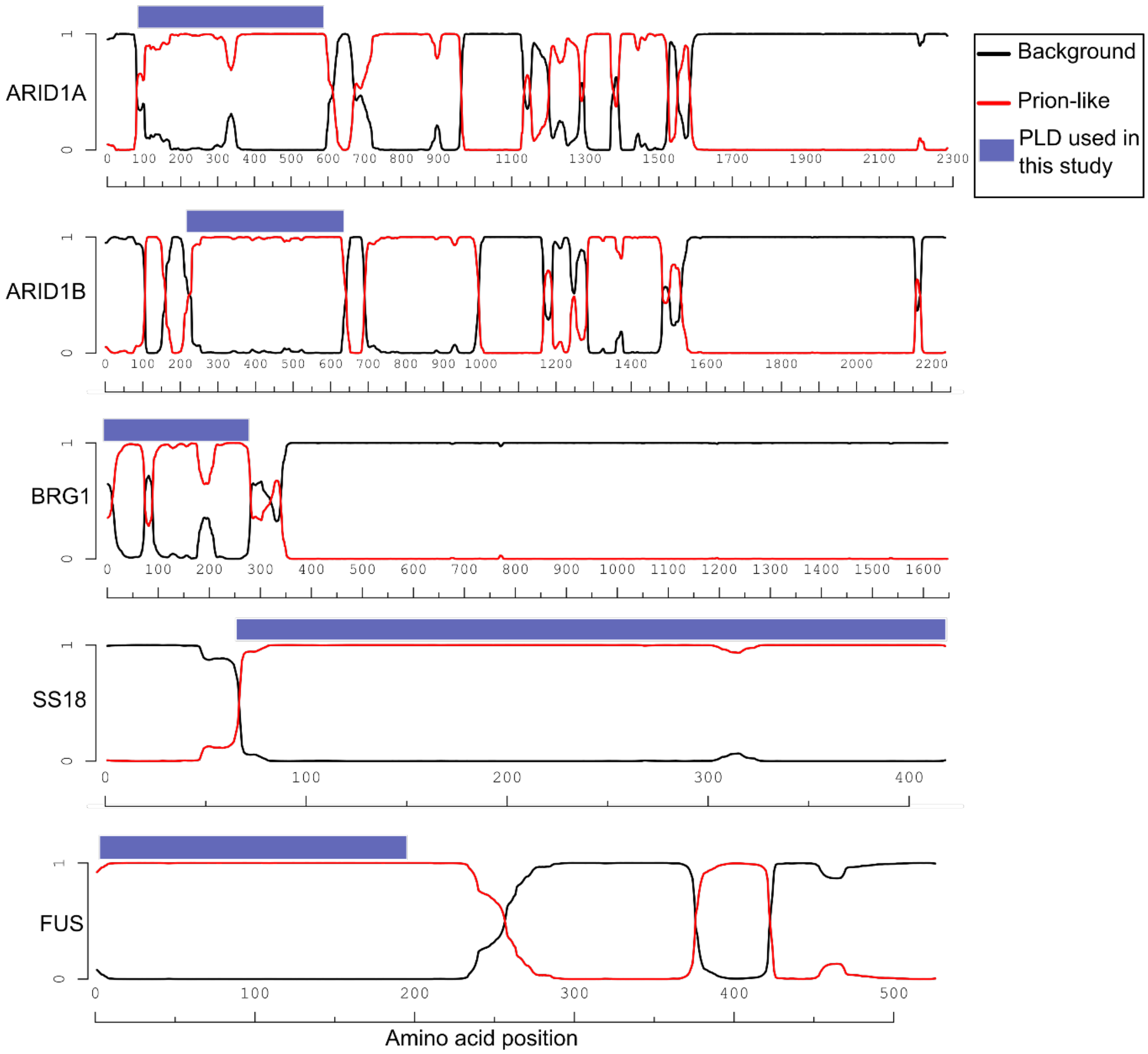


**Figure S1:** PLAAC analysis [7] showing regions with high prion-propensity for the four subunits of mSWI/SNF complex (ARID1A, ARID1B, BRG1, and SS18) and FUS. Domains corresponding to blue bars were used in this study.

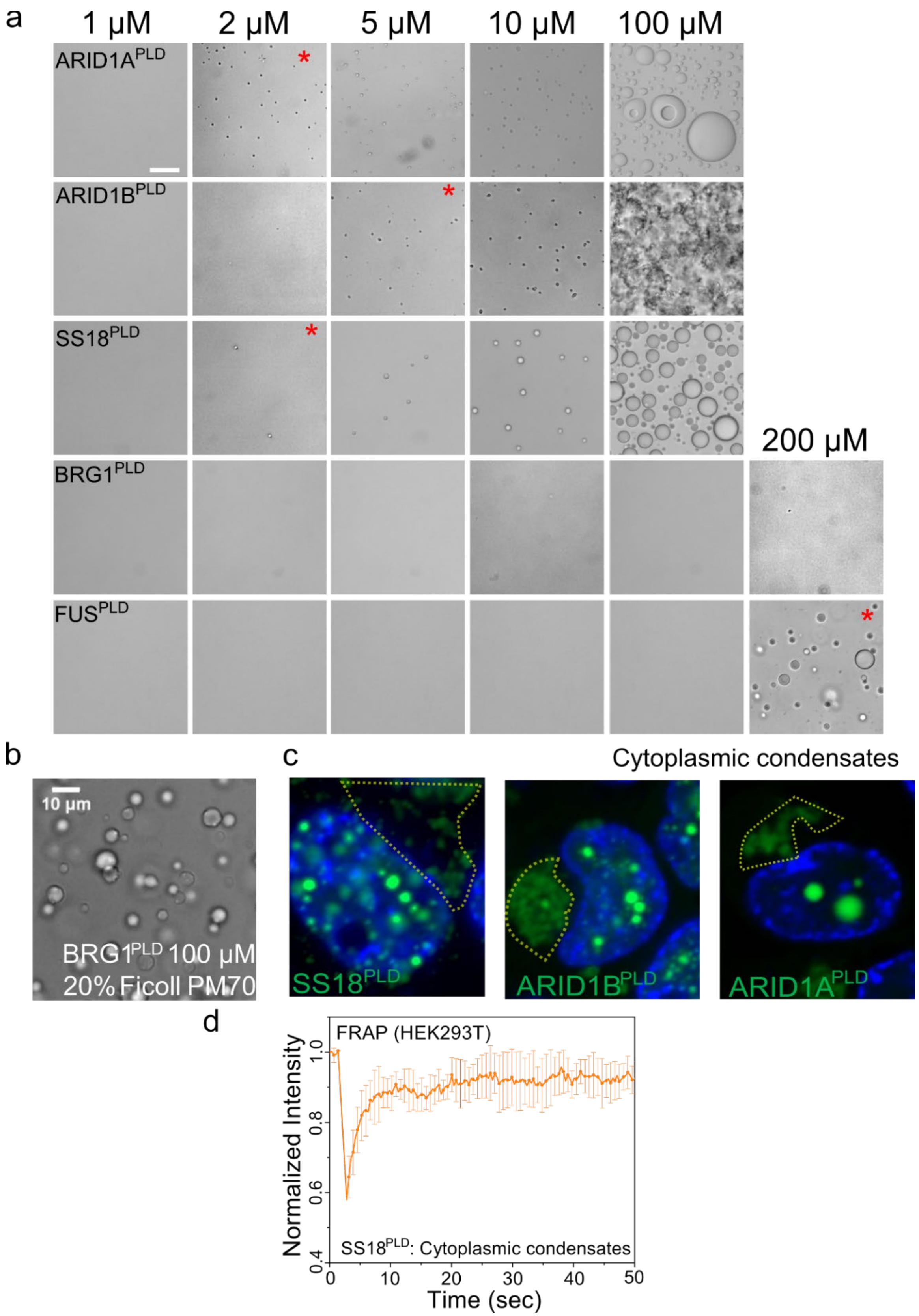
a
1 μM
2 μM
5 μM
10 μM
100 μM
200 μM
ARID1A^PLD
ARID1B^PLD
SS18^PLD
BRG1^PLD
FUS^PLD
b
10 μm
BRG1^PLD 100 μM
20% Ficoll PM70
c
Cytoplasmic condensates
SS18^PLD
ARID1B^PLD
ARID1A^PLD
d
FRAP (HEK293T)
Normalized Intensity
SS18^PLD: Cytoplasmic condensates
Time (sec)
0.4
0.6
0.8
1.0
0
10
20
30
40
50

**Figure S2: a)** Differential interference contrast (DIC) microscopy images showing concentration titrations of recombinantly purified $FUS^{PLD}$ and mSWI/SNF PLDs ($ARID1A^{PLD}$, $ARID1B^{PLD}$, $SS18^{PLD}$, and $BRG1^{PLD}$). The red asterisk indicates the concentration at which condensates were first observed. The scale bar is 10 μm. **b)** Phase separation of $BRG1^{PLD}$ was observed at 100 μM protein concentration with 20% Ficoll PM70. **c)** Irregular-shaped cytoplasmic condensates of GFP-labeled PLDs in HEK293T cells are displayed within the yellow dashed lines. Hoechst was used to stain the cell nucleus, which is shown in blue. **d)** FRAP curve for cytoplasmic condensates of GFP-tagged $SS18^{PLD}$ in HEK293T cells. The average intensity and standard deviation of the intensity profiles are shown as a function of time (n = 3).

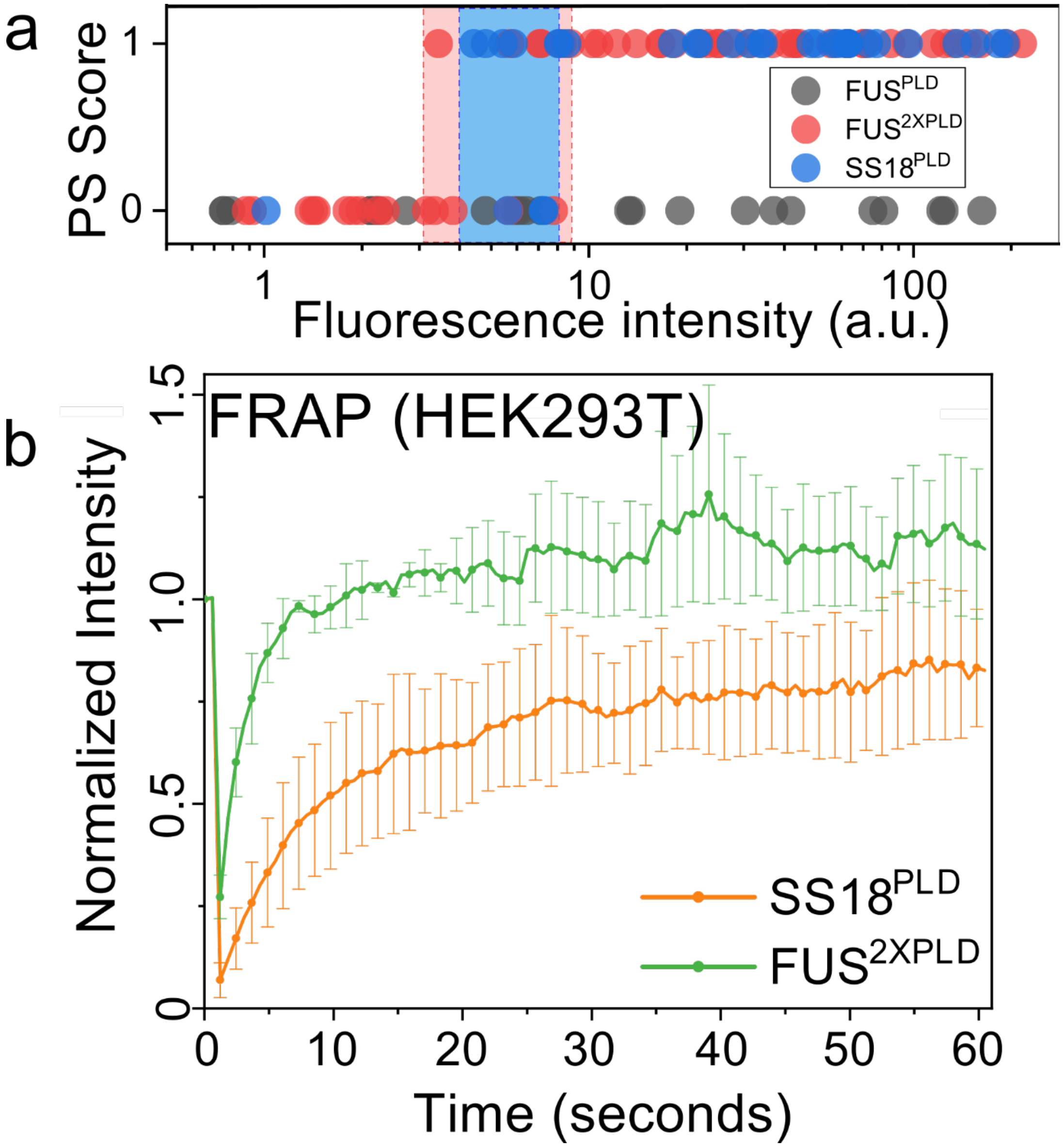


**Figure S3: a)** The phase separation capacity is quantified over various levels of nuclear protein concentration for FUS$^{PLD}$, FUS$^{2XPLD}$ and SS18$^{PLD}$ and presented as a state diagram. A phase separation (PS) score of '1' indicates the presence of nuclear condensates and a PS score of '0' represents diffused expression patterns. The shaded regions represent the transition concentrations (n = total of 28-55 cells from two replicates). **b)** FRAP curve for condensates of GFP-tagged FUS$^{2XPLD}$ and SS18$^{PLD}$ in HEK293T cells. The average intensity and standard deviation of the intensity profiles are shown over time (n = 3). The scale bar is 5 μm for all images.

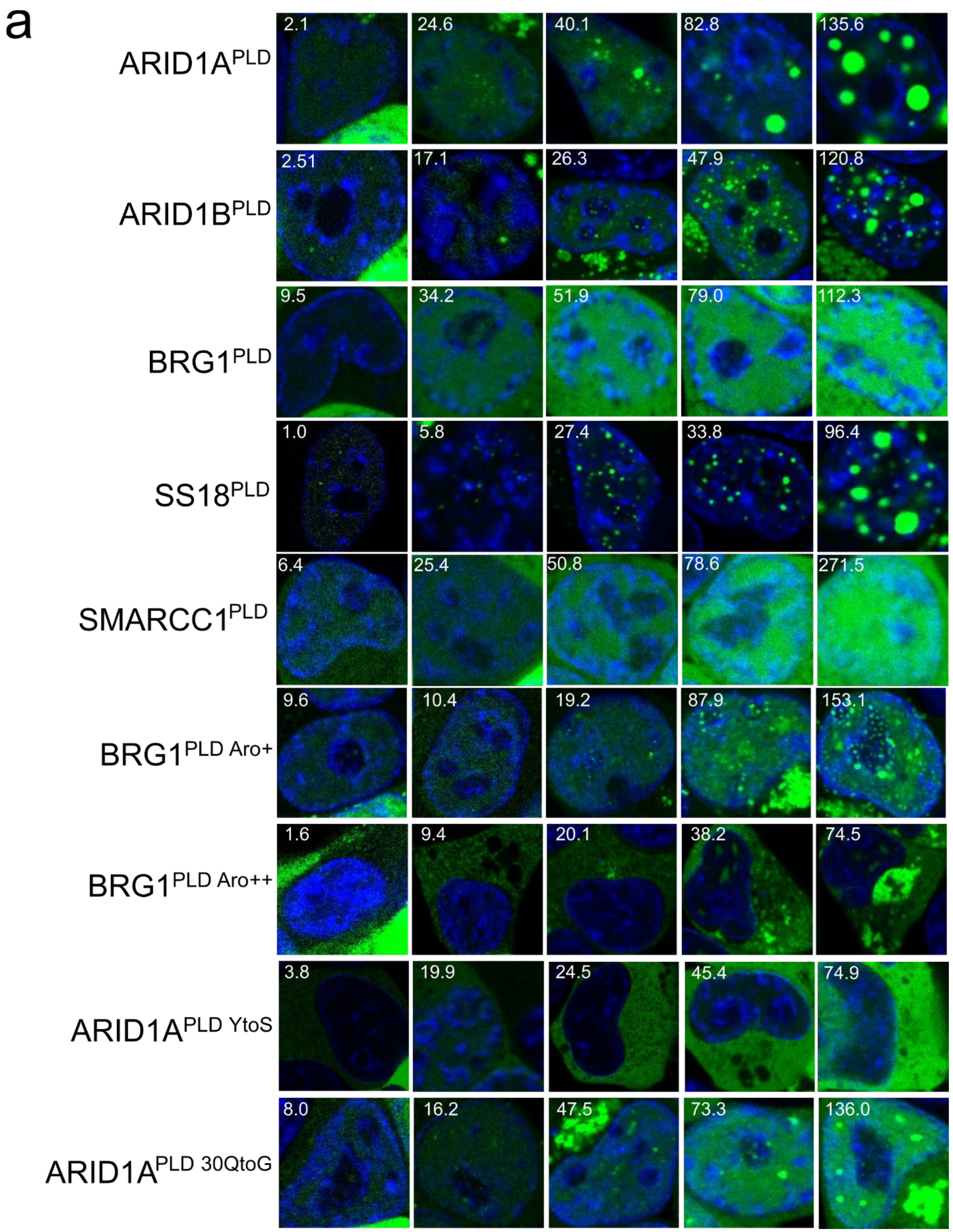

a
ARID1A^PLD
2.1
24.6
40.1
82.8
135.6
ARID1B^PLD
2.51
17.1
26.3
47.9
120.8
BRG1^PLD
9.5
34.2
51.9
79.0
112.3
SS18^PLD
1.0
5.8
27.4
33.8
96.4
SMARCC1^PLD
6.4
25.4
50.8
78.6
271.5
BRG1^PLD Aro+
9.6
10.4
19.2
87.9
153.1
BRG1^PLD Aro++
1.6
9.4
20.1
38.2
74.5
ARID1A^PLD YtoS
3.8
19.9
24.5
45.4
74.9
ARID1A^PLD 30QtoG
8.0
16.2
47.5
73.3
136.0

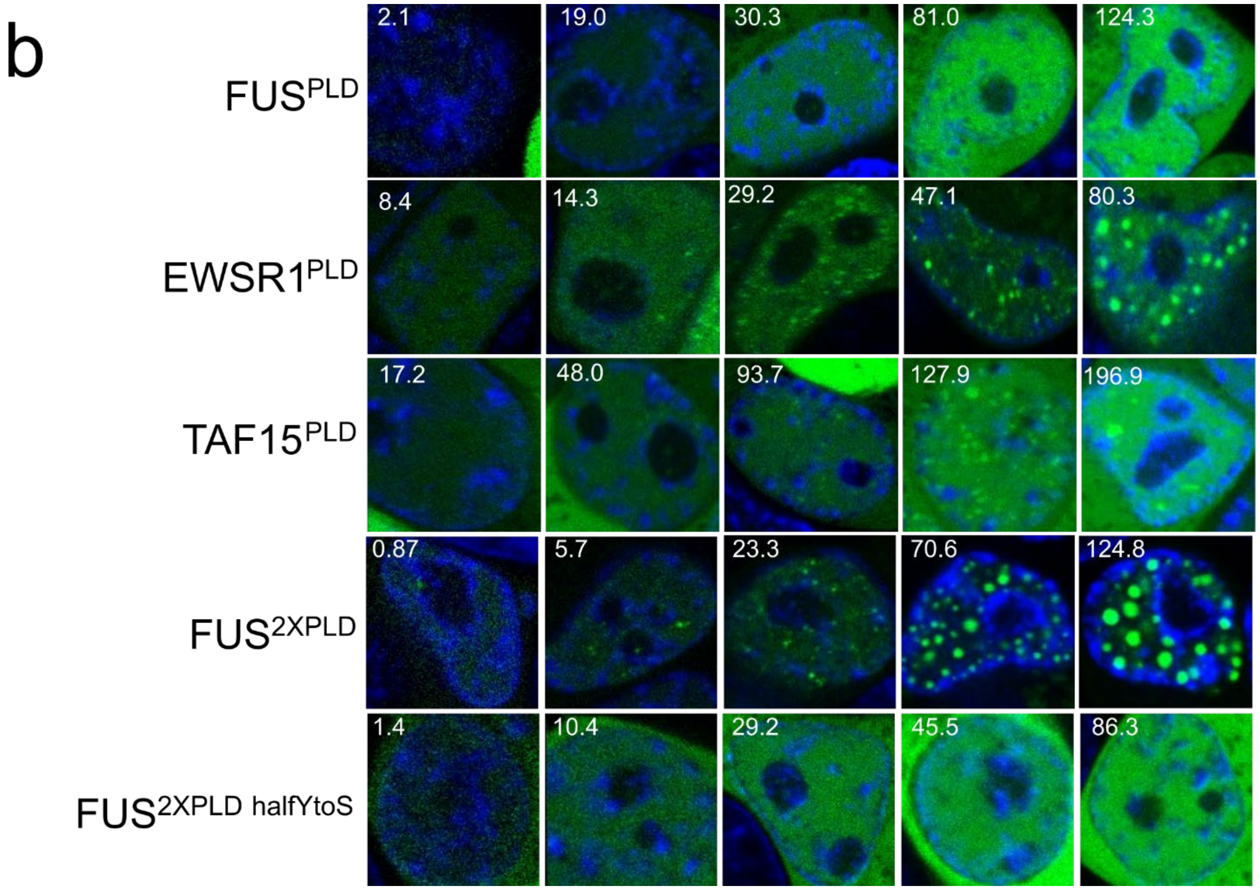


**Figure S4:** Fluorescence microscopy images of HEK293T cells expressing GFP-tagged PLDs and variants of **a)** mSWI/SNF subunits, and **b)** FET proteins at varying expression levels. The mean GFP intensity values are noted in each image. Hoechst was used to stain the cell nucleus, which is shown in blue. For BRG1$^{PLD\ Aro++}$ and ARID1A$^{PLD\ YtoS}$, mean cytoplasmic intensities are noted since their expression was predominantly cytoplasmic. For some mSWI/SNF subunit PLDs, higher expression level has been noted to result in the formation of irregular cytoplasmic condensates.

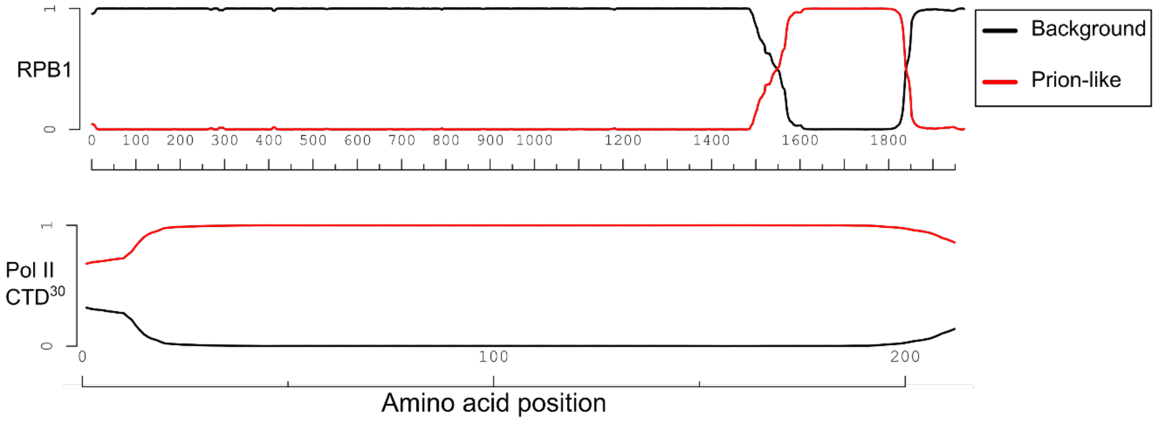


**Figure S5:** PLAAC analysis [7] showing regions with high prion-propensity for the human RNA Polymerase II subunit RPB1 (*Top panel*) and the first 30 repeats of the heptad (Pol II CTD [30]) used in this study (*bottom panel*).

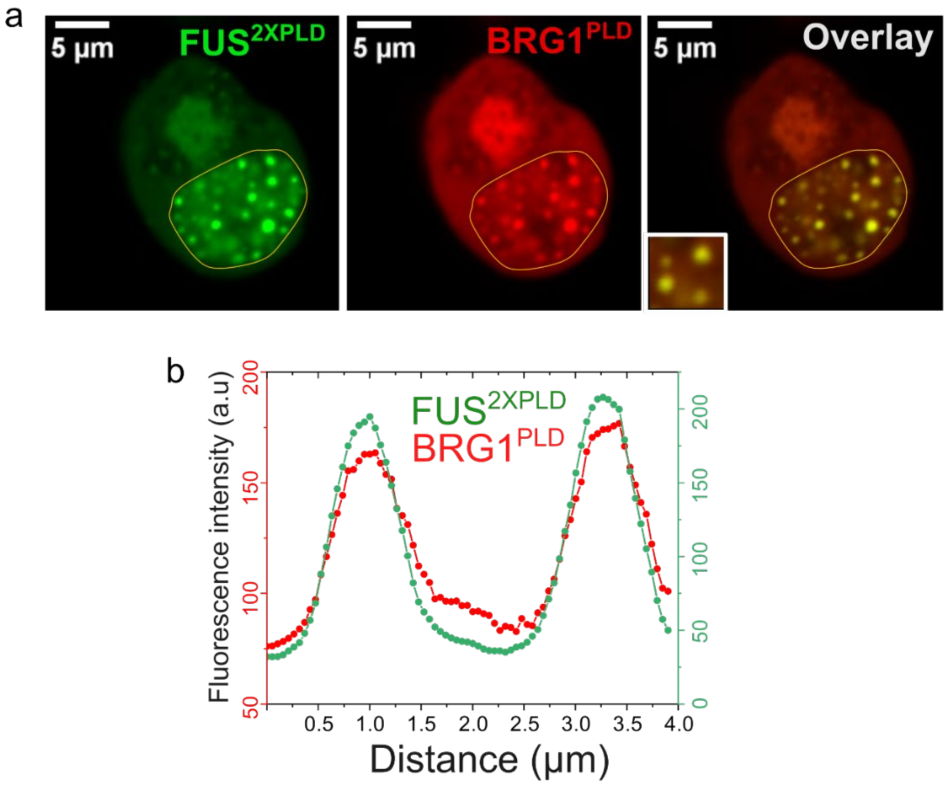


**Figure S6: a)** HEK293T cells co-expressing GFP-FUS$^{2XPLD}$ and mCherry-BRG1$^{PLD}$. **b)** The degree of colocalization is displayed as intensity profiles for condensates shown in the inset image. Green represents the intensity profile of GFP-FUS$^{2XPLD}$ and red represents the intensity profile for mCherry-BRG1$^{PLD}$.

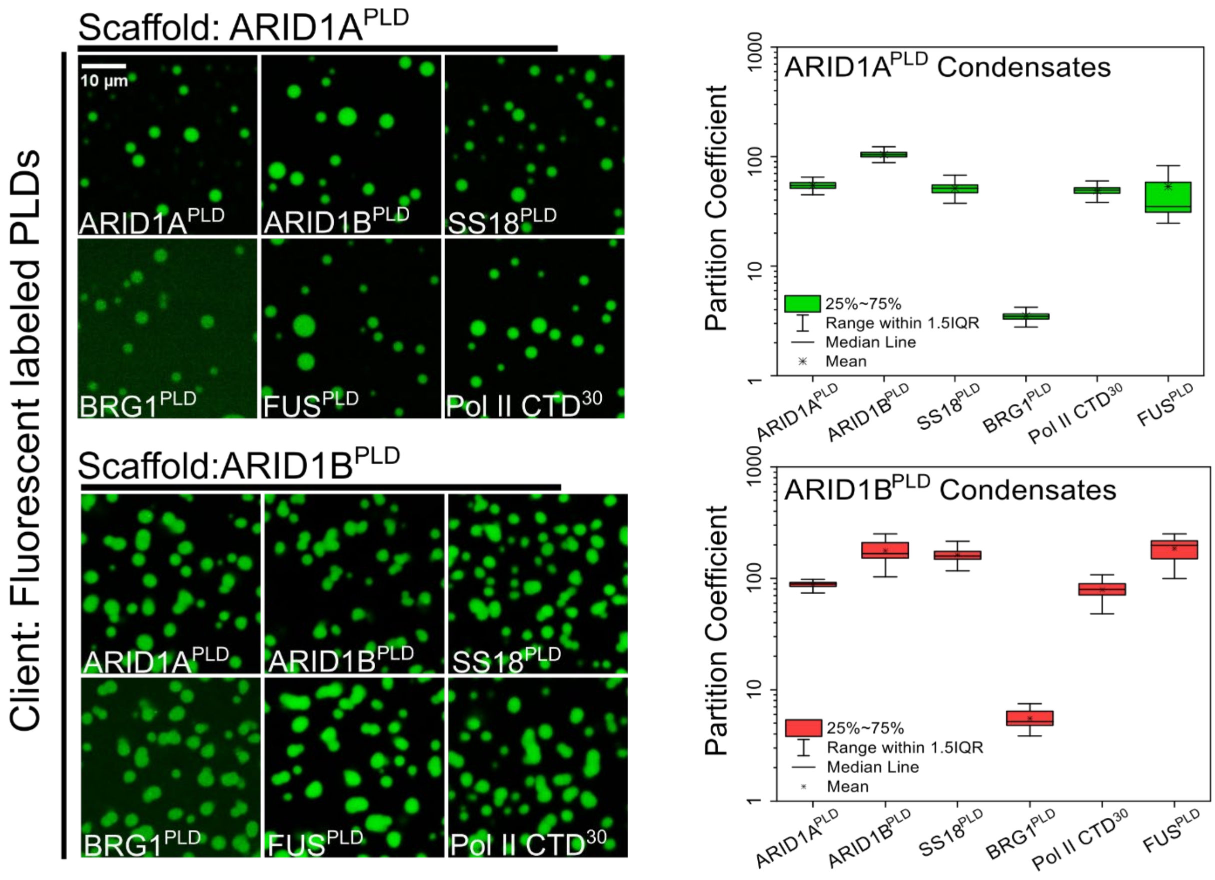


**Figure S7:** Partitioning of AlexaFluor488 labeled client PLDs (ARID1A$^{PLD}$, ARID1B$^{PLD}$, SS18$^{PLD}$, BRG1$^{PLD}$, RNA Polymerase II CTD$^{30}$, and FUS$^{PLD}$) within condensates of ARID1A$^{PLD}$ and ARID1B$^{PLD}$. Enrichment (partition coefficient) is calculated as shown in Fig. 3 in the main text and displayed as a box-and-whisker plot.

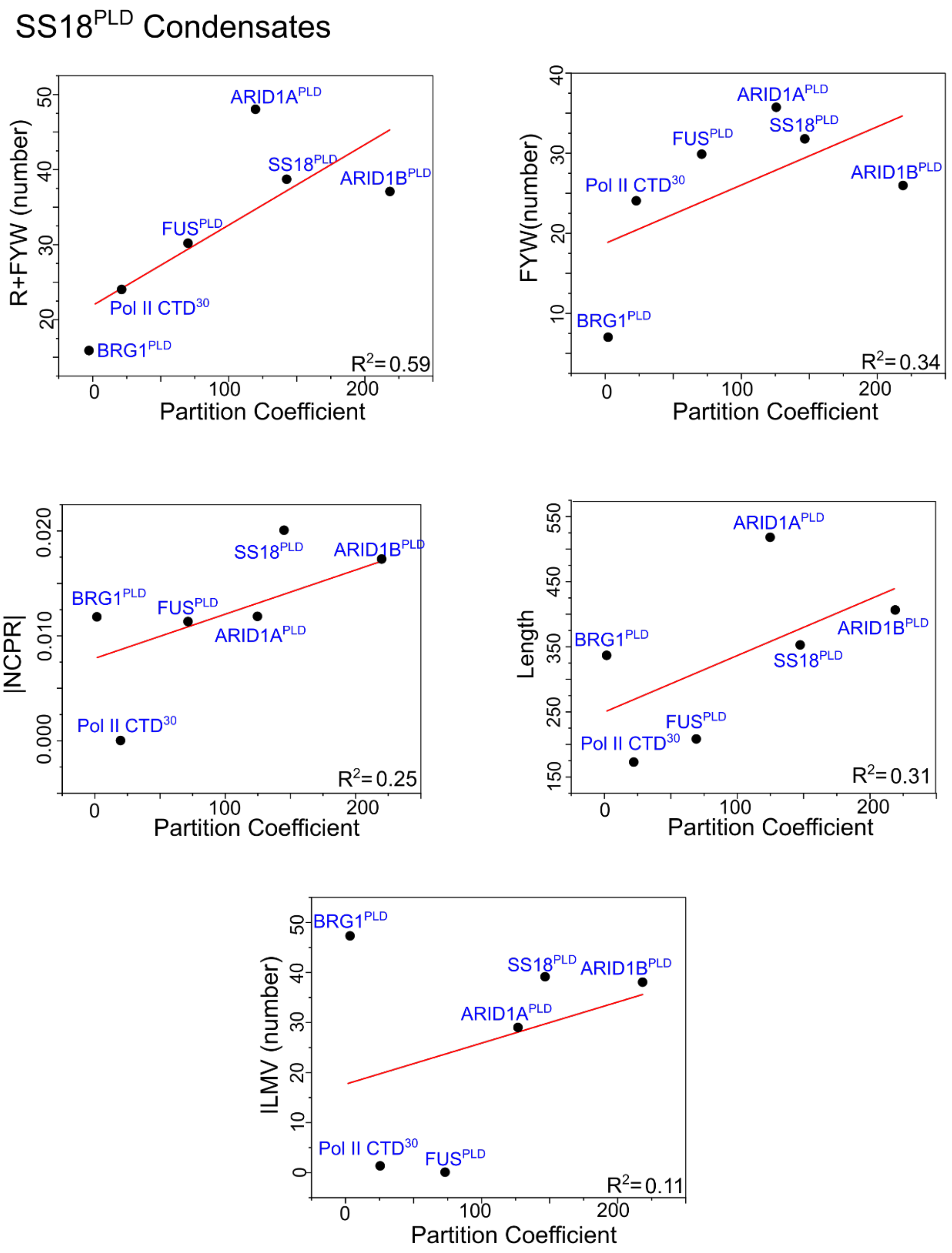


**Figure S8:** Linear regression analysis of in vitro partition coefficients in condensates formed by 50 µM SS18$^{PLD}$ against various sequence features of the client PLDs (ARID1A$^{PLD}$, SS18$^{PLD}$, ARID1B$^{PLD}$, FUS$^{PLD}$, Pol II CTD$^{30}$, and BRG1$^{PLD}$). Sequence features included in the analysis are FYW: phenylalanine, tyrosine, and tryptophan residues representing the aromatic side chains; R: arginine; NCPR: net charge per residue; Chain length; ILMV: isoleucine, leucine, methionine, and valine representing the hydrophobic residues.

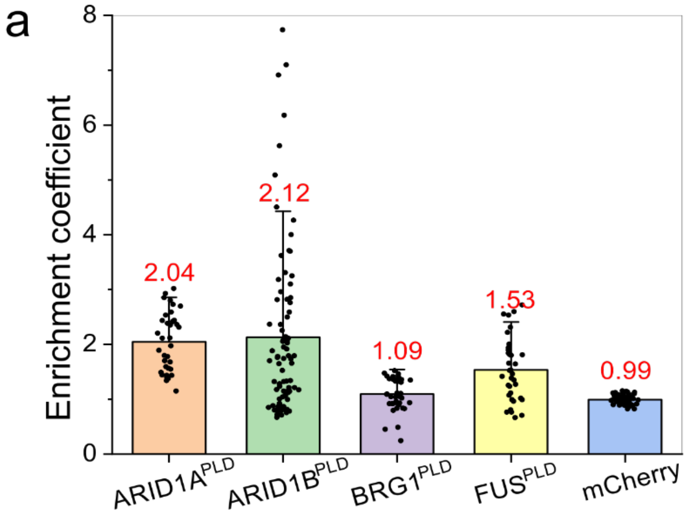


| *p*-value | **ARID1B$^{PLD}$** | **BRG1$^{PLD}$** | **FUS$^{PLD}$** | **mCherry** |
|---|---|---|---|---|
| **ARID1A$^{PLD}$** | 7.49E-01 | 3.49E-15 | 1.56E-04 | 2.62E-29 |
| **ARID1B$^{PLD}$** | | 5.87E-05 | 2.48E-02 | 1.12E-08 |
| **BRG1$^{PLD}$** | | | 9.80E-05 | 5.58E-04 |
| **FUS$^{PLD}$** | | | | 2.00E-11 |

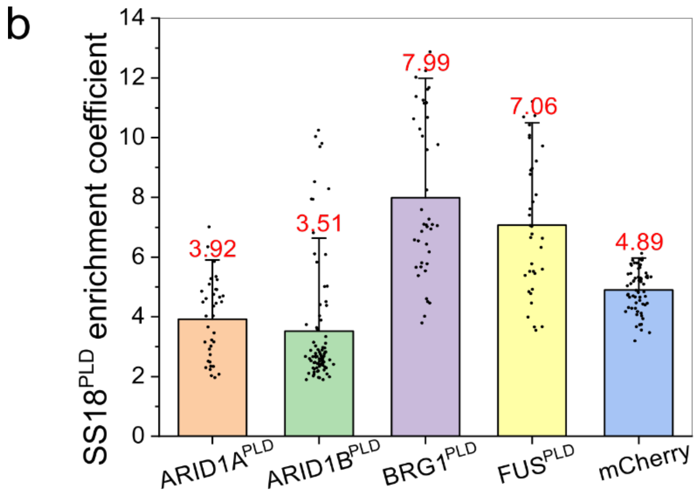


**Figure S9: a)** Enrichment coefficients of mCherry-tagged PLDs (ARID1A$^{PLD}$, ARID1B$^{PLD}$, BRG1$^{PLD}$, FUS$^{PLD}$, and mCherry alone) within condensates formed by GFP-SS18$^{PLD}$ in HEK293T cells. Enrichment is calculated as the ratio of mean intensities from the dense phase and the dilute phase. The average enrichment coefficient score is shown in red for each PLD and the mCherry control. (n = 36-85 condensates from 5-8 cells). Student's t-test was used to calculate significance for each of the constructs and the p-values are tabulated. **b)** Self-partitioning of GFP-SS18$^{PLD}$

within condensates formed when co-expressed with different mCherry-tagged client PLDs as indicated.

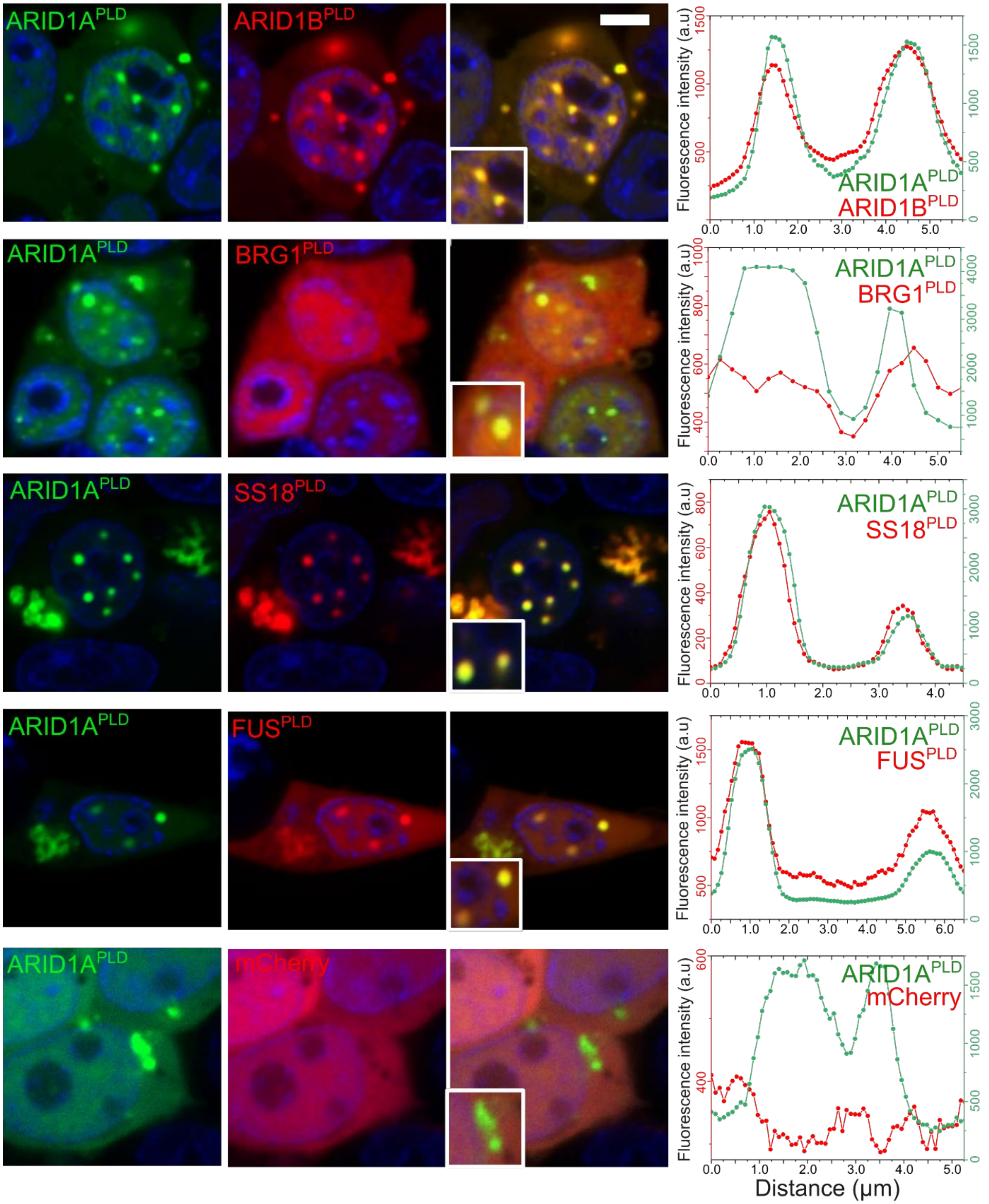


**Figure S10:** HEK293T cells co-expressing GFP-ARID1A$^{PLD}$ and either one of the mCherry-tagged PLDs (SS18$^{PLD}$, ARID1B$^{PLD}$, FUS$^{PLD}$, and BRG1$^{PLD}$) or mCherry alone. The degree of co-localization is displayed as intensity profiles for condensates shown in the inset images. Green represents the intensity profile of GFP-ARID1A$^{PLD}$ and red represents the intensity profile for mCherry-PLD. The scale bar is 10 μm.

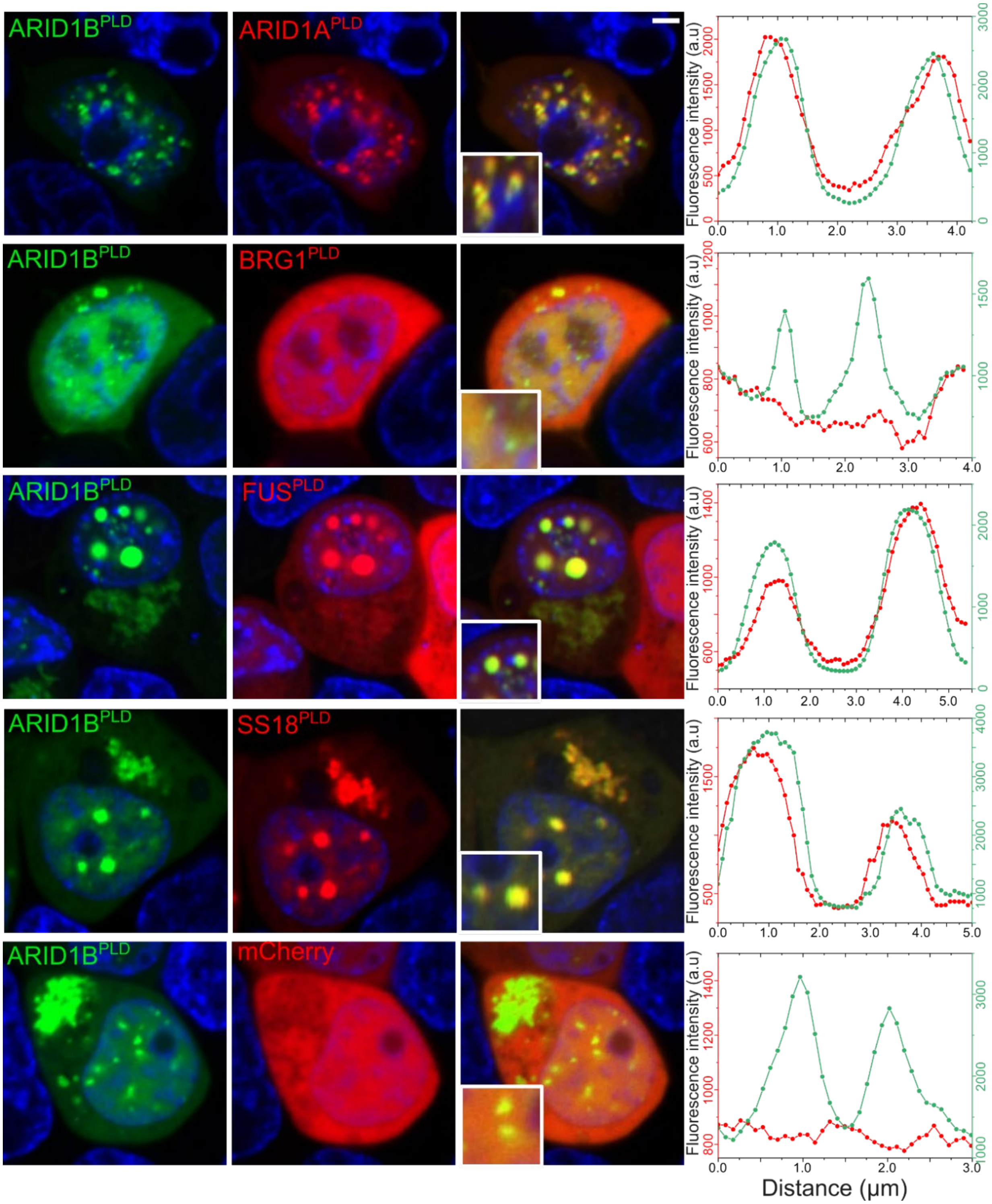


**Figure S11:** HEK293T cells co-expressing GFP-ARID1B$^{PLD}$ and either one of the mCherry-tagged PLDs (SS18$^{PLD}$, ARID1B$^{PLD}$, FUS$^{PLD}$, and BRG1$^{PLD}$) or mCherry alone. The degree of colocalization is displayed as intensity profiles for condensates shown in the inset images. Green represents the intensity profile of GFP-ARID1A$^{PLD}$ and red represents the intensity profile for mCherry-PLD. The scale bar is 5 μm.

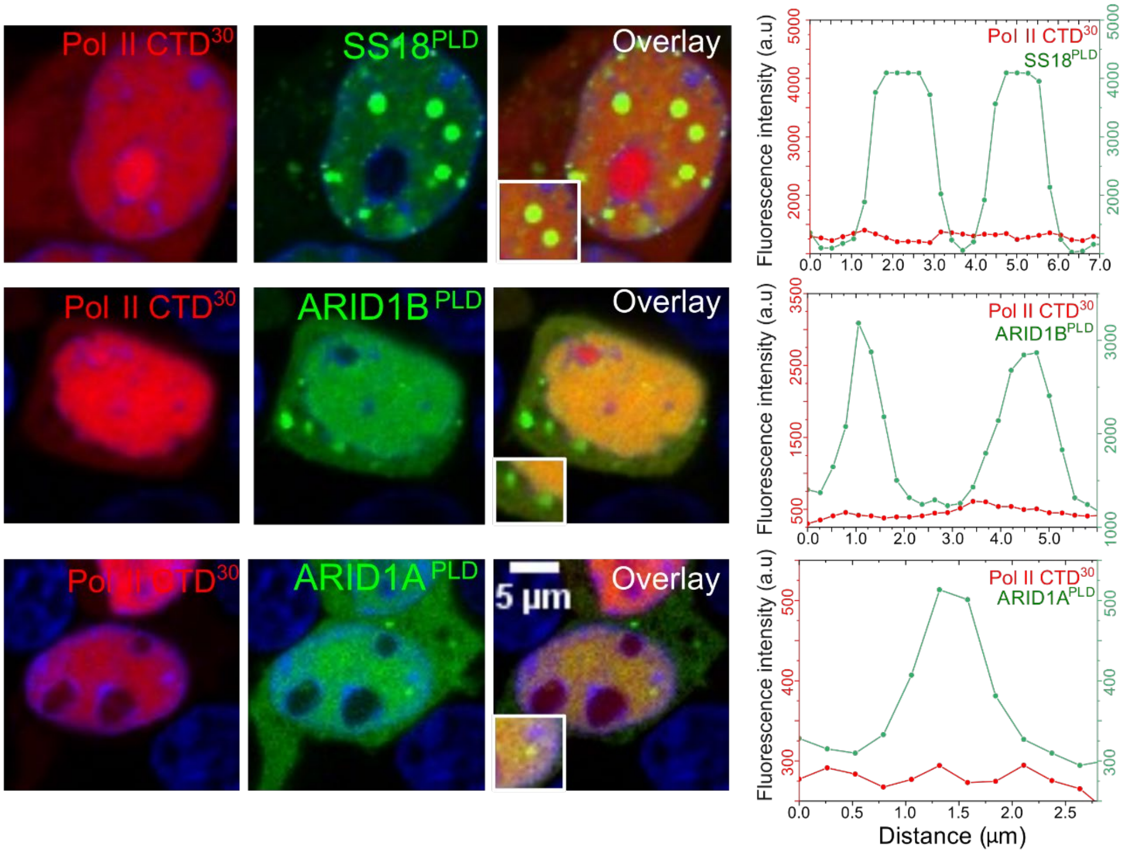


**Figure S12:** HEK293T cells co-expressing mCherry-RNA Pol II CTD$^{30}$ and either one of the GFP tagged PLDs (ARID1A$^{PLD}$, SS18$^{PLD}$, ARID1B$^{PLD}$). The intensity profile is shown for condensates in the inset images. The degree of colocalization is displayed as intensity profiles for condensates present in the inset images. Green represents the intensity profile of GFP-PLDs and red represents the intensity profile for mCherry-Pol II CTD$^{30}$.

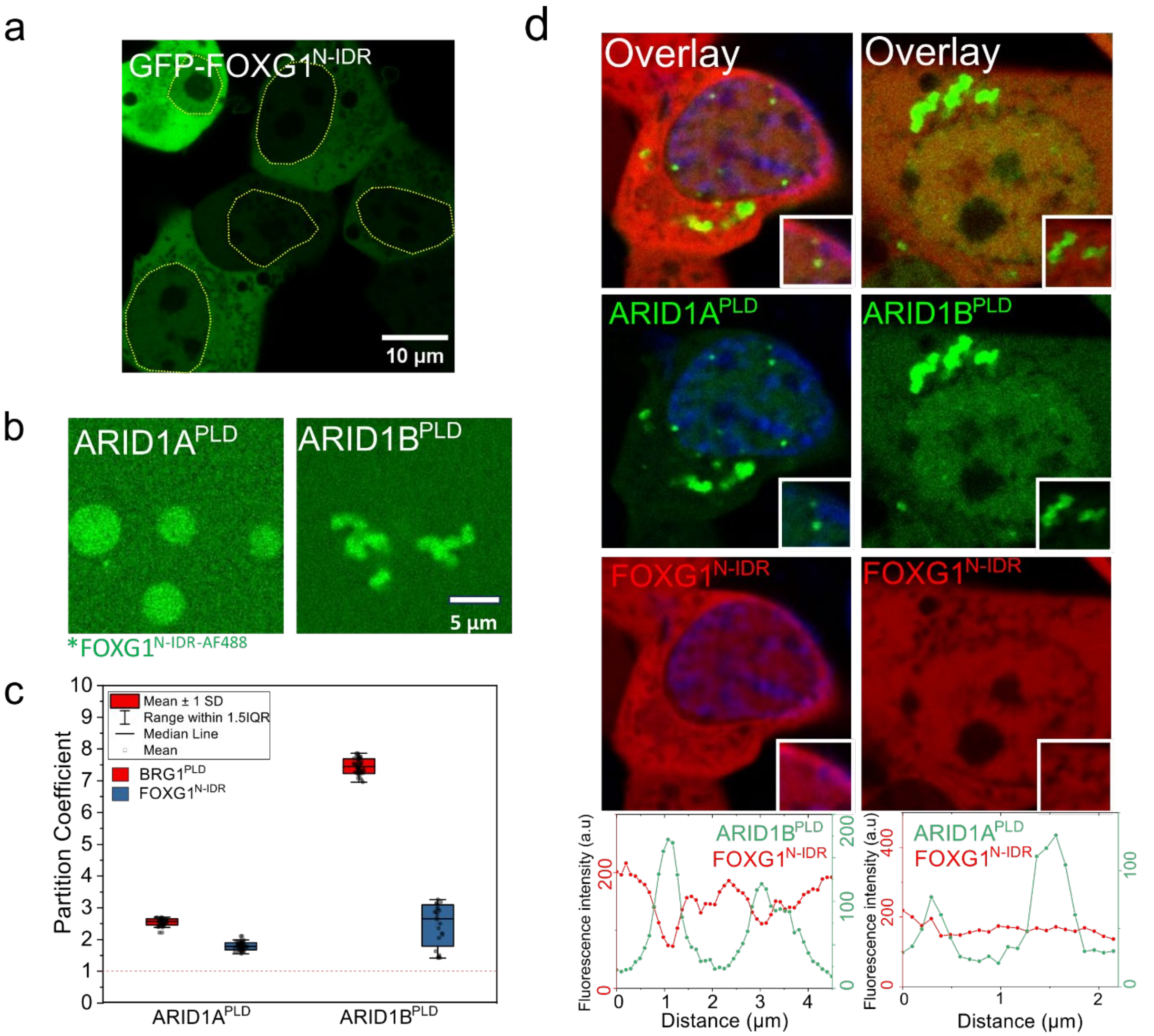


**Figure S13: a)** Fluorescence microscopy image of HEK293T cells expressing GFP-tagged FOXG1$^{N\text{-}IDR}$. Yellow dashed lines indicate the nuclear periphery. **b)** Partitioning of AlexaFluor488 labeled FOXG1$^{N\text{-}IDR}$ within condensates formed by ARID1A$^{PLD}$ and ARID1B$^{PLD}$ (50 µM), respectively. **c)** Enrichment is calculated as partition coefficient and displayed as a box-and-whisker plot for both FOXG1$^{N\text{-}IDR}$ and BRG1$^{PLD}$ within these condensates (n = total of 25-50 droplets). **d)** HEK293T cells co-expressing GFP-ARID1A$^{PLD}$ or GFP-ARID1B$^{PLD}$ and mCherry-tagged FOXG1$^{N\text{-}IDR}$.The degree of colocalization is displayed as intensity profiles for condensates shown in the inset images. Green represents the intensity profile of the GFP-tagged construct and red represents the profile for mCherry-tagged construct.

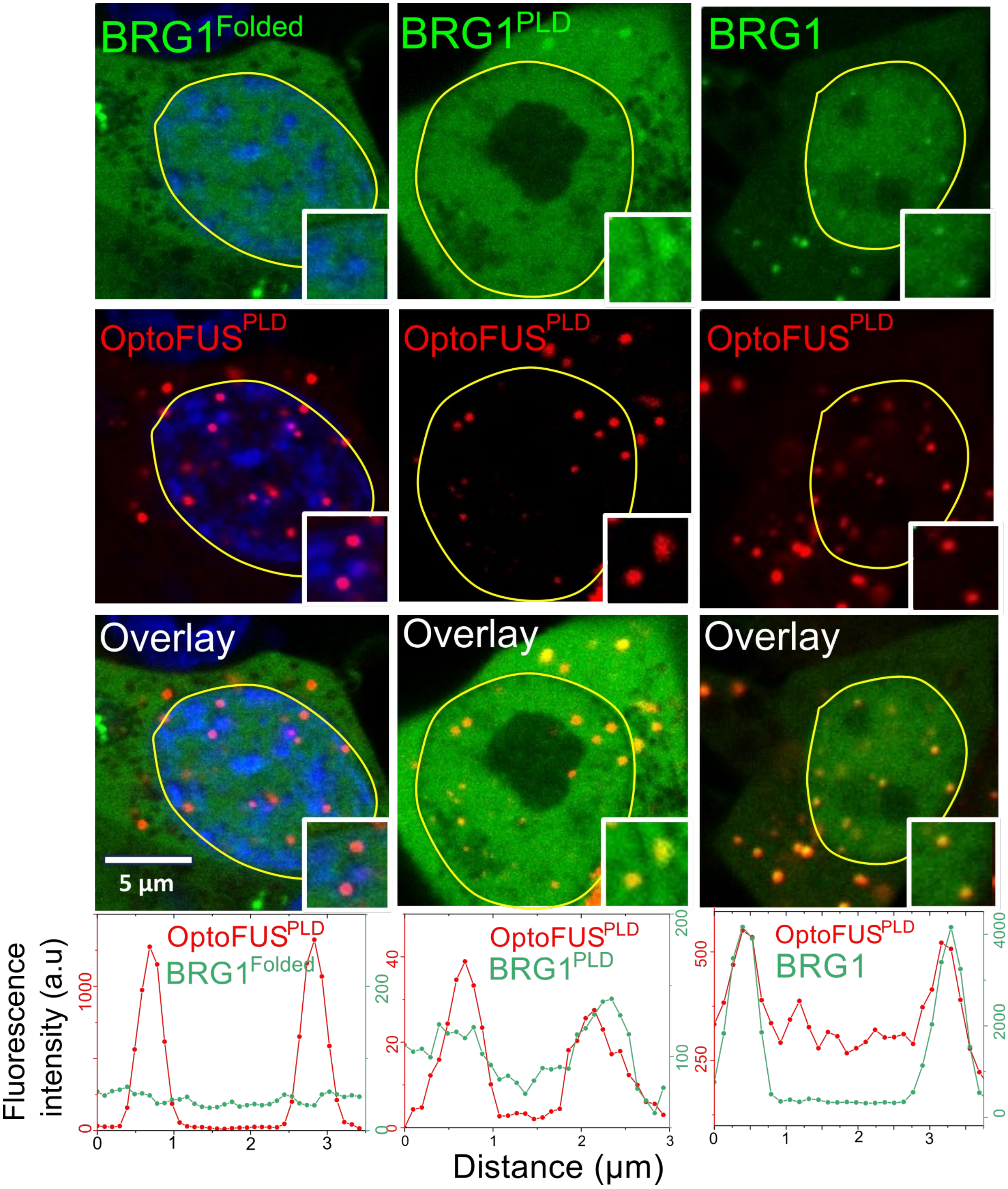


**Figure S14:** HEK293T cells co-expressing BRG1$^{PLD}$, BRG1$^{Folded}$ or full-length BRG1 and mCherry-tagged OptoFUS$^{PLD}$ construct. The degree of colocalization is displayed as intensity profiles for condensates shown in the inset images. Green represents the intensity profile of the GFP-tagged construct and red represents the profile for mCherry-tagged construct. Yellow lines indicate the nuclear periphery.

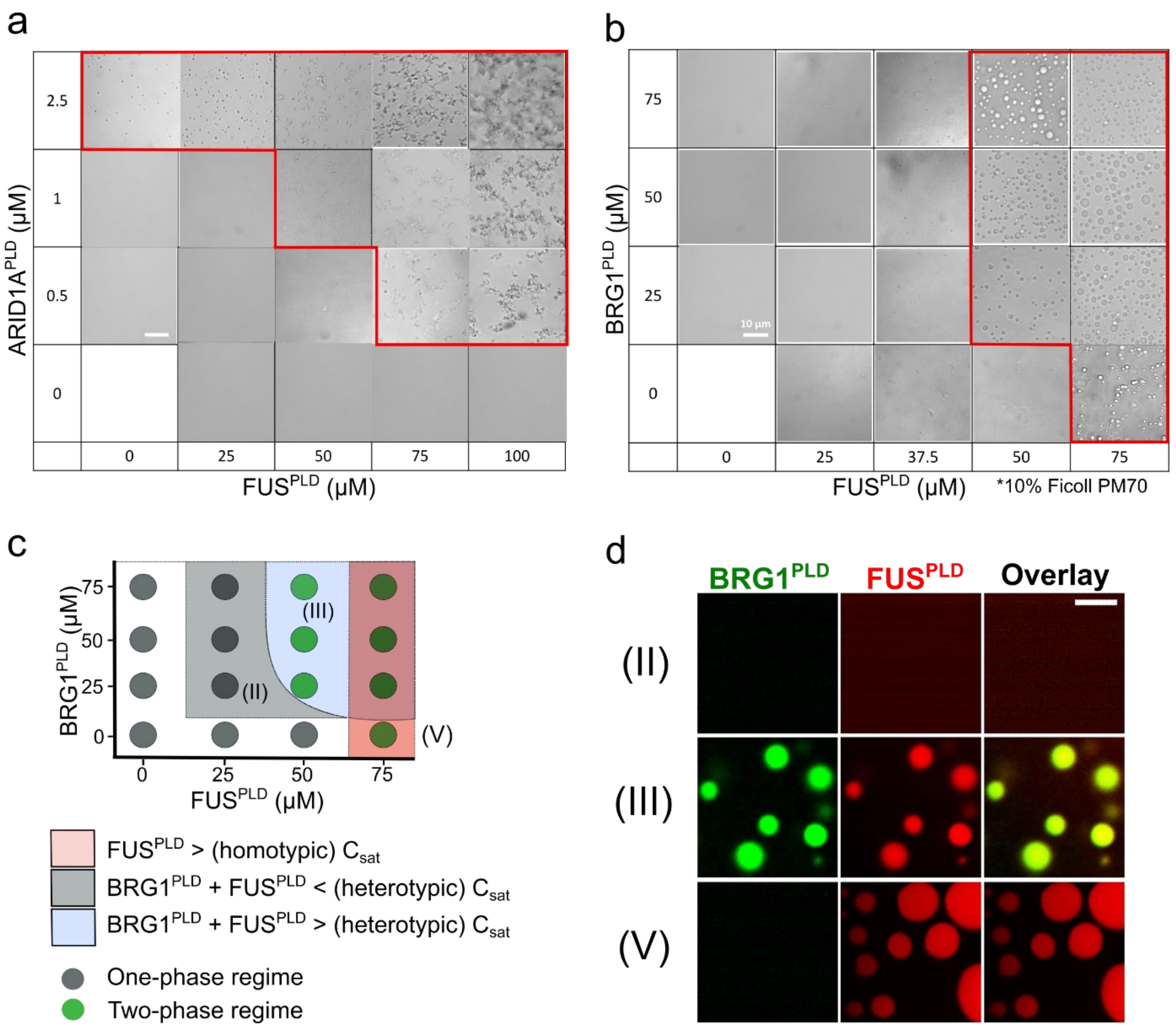


**Figure S15:** DIC images of the heterotypic PLD mixtures containing FUS$^{PLD}$ and either **a)** ARID1A$^{PLD}$ or **b)** BRG1$^{PLD}$ at indicated concentrations. The images within the red-boxed region represent the two-phase regime. The FUS$^{PLD}$ and BRG1$^{PLD}$ ternary mixture contained 10% Ficoll PM70 in the buffer. **c)** Co-phase diagram of FUS$^{PLD}$ and BRG1$^{PLD}$ showing a decrease in saturation concentration of heterotypic PLD mixtures (regime III). The green circles indicate a two-phase regime, and the grey circles indicate a single-phase regime. The legend describes the shaded regions. **d)** Fluorescence microscopy images of samples from the indicated regions on the phase diagram in (**c**). These images show the formation of monophasic co-condensates of FUS$^{PLD}$ and BRG1$^{PLD}$. BRG1$^{PLD}$ is labeled with AlexaFluor488 and FUS$^{PLD}$ is labeled with AlexaFluor594. The scale bar is 5 microns.

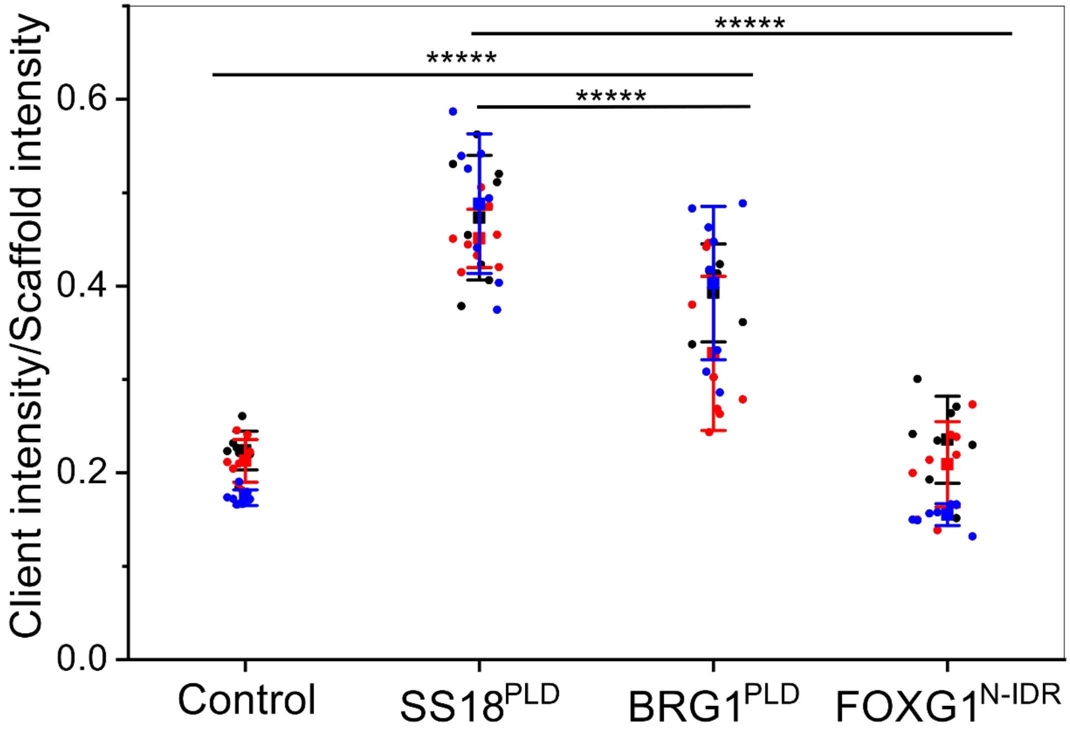


**Figure S16:** Statistical analysis for the bead halo assay data shown in Fig. 6a-c in the main text. Three trials were combined (with points for each of the trials shown in different colors) to calculate the significance using Student’s t-test. ***** P ≤ 0.00001. Here the control refers to the AlexaFluor488-labeled His$_6$-MBP containing a short linker peptide (GGGCGGG) without any PLDs.

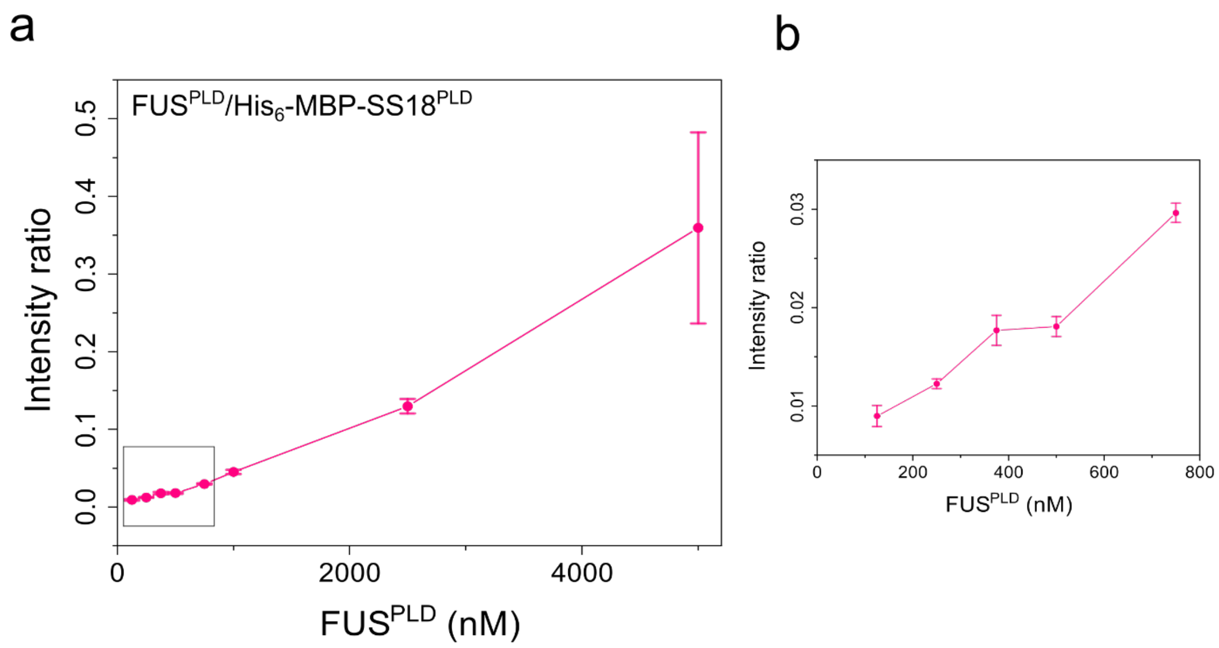


**Figure S17: a)** 250 nM of AlexaFluor488 labeled His$_6$-MBP-SS18$^{PLD}$ was attached to Ni-NTA beads. AlexaFluor594 labeled FUS$^{PLD}$ was titrated from 125 nM to 5 µM. Binding was quantified using the ratio of the fluorescence intensity (fluorescence signal from FUS$^{PLD}$/fluorescence signal from SS18$^{PLD}$) on the surface of the beads and plotted as a line plot with mean and standard deviation (n = 8 beads). **b)** The region within the square box in (**a**) is enlarged and displayed here.

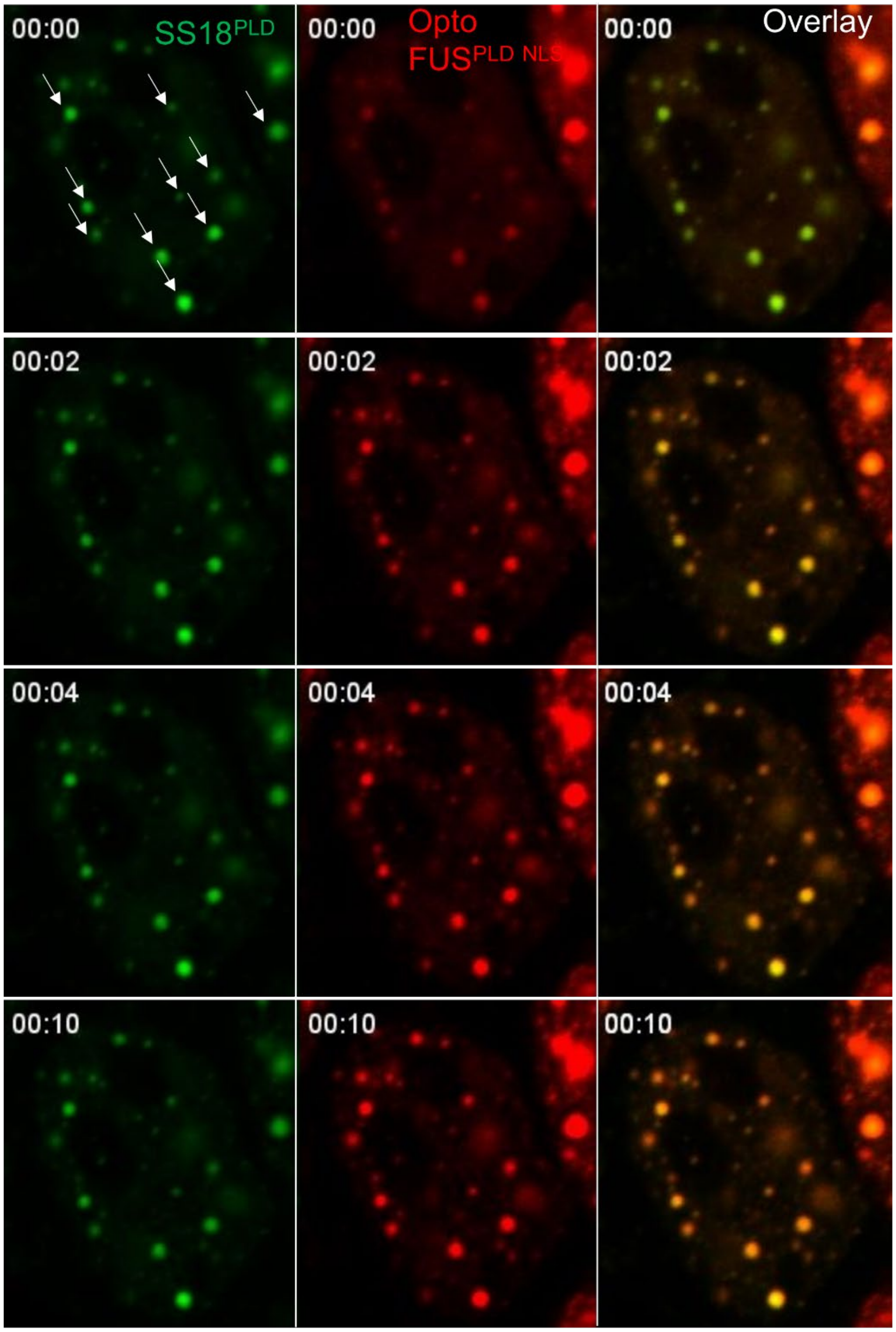


**Figure S18.** HEK293T cells co-expressing OptoFUS$^{PLD-NLS}$ (Cry2-mCherry-FUS$^{PLD-NLS}$) and GFP-SS18$^{PLD}$. Pre-existing GFP-SS18$^{PLD}$ condensates, which are indicated by white arrows, serve as seeds to nucleate OptoFUS$^{PLD-NLS}$ condensates upon blue light activation. The corresponding video is shown as Movie S3.

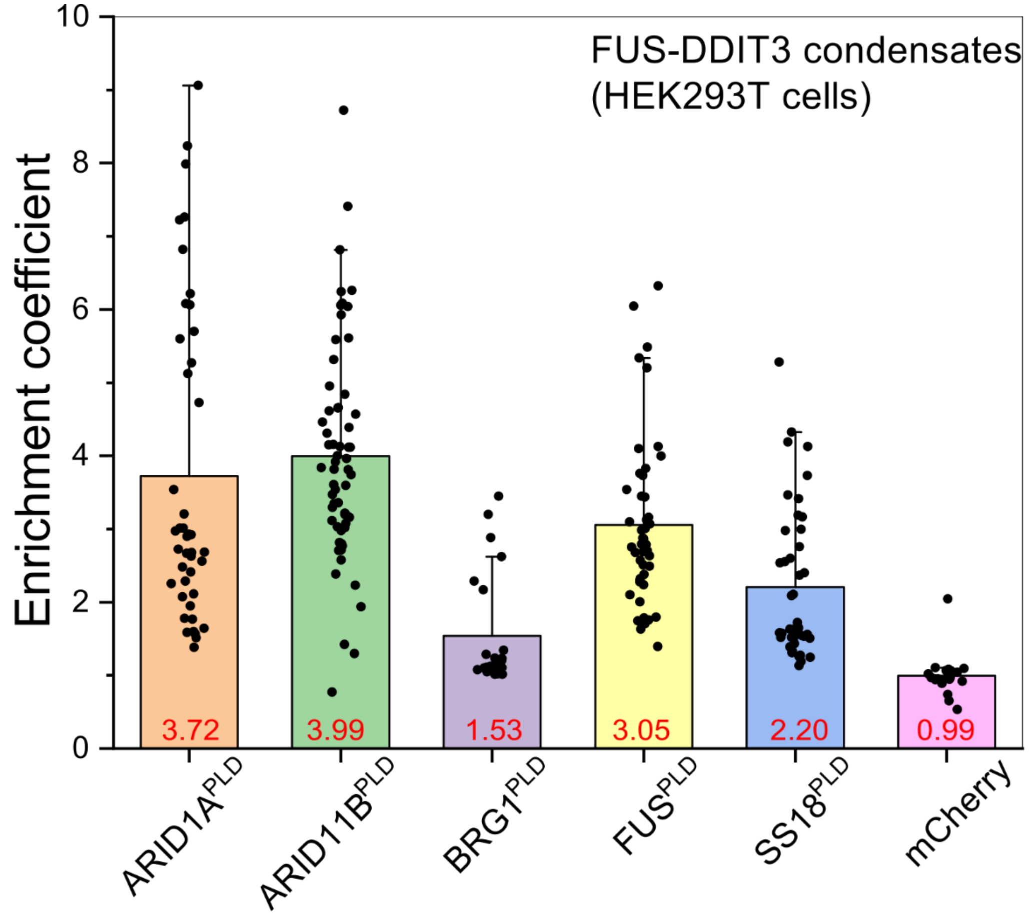


| *p*-value | ARID1B$^{PLD}$ | BRG1$^{PLD}$ | FUS$^{PLD}$ | SS18$^{PLD}$ | mCherry |
|---|---|---|---|---|---|
| **ARID1A$^{PLD}$** | 4.46E-01 | 9.72E-06 | 6.85E-02 | 6.03E-05 | 1.56E-07 |
| **ARID1B$^{PLD}$** | | 2.69E-11 | 5.62E-04 | 4.35E-10 | 1.47E-14 |
| **BRG1$^{PLD}$** | | | 2.11E-07 | 6.77E-03 | 3.07E-03 |
| **FUS$^{PLD}$** | | | | 4.01E-04 | 1.23E-11 |
| **SS18$^{PLD}$** | | | | | 9.17E-07 |

**Figure S19:** Enrichment coefficients of mCherry-tagged PLDs (ARID1A$^{PLD}$, ARID1B$^{PLD}$, BRG1$^{PLD}$, FUS$^{PLD}$, SS18$^{PLD}$ and mCherry alone) within condensates formed by GFP-FUS-DDIT3 in HEK293T cells. Enrichment is calculated as the ratio of mean intensities from the dense phase and the dilute phase. The average enrichment coefficient score is shown in red for each client. (n = 25-65 condensates from 4-8 cells). Student's t-test was used to calculate significance for each of the constructs and the p-values are tabulated.

### Supplementary Movies

**Movie S1:** HEK293T cells co-expressing OptoFUS$^{PLD\text{-}NLS}$ (Cry2-mCherry-FUS$^{PLD\text{-}NLS}$) and GFP-SS18$^{PLD}$ below their saturation concentrations. Upon blue light activation, OptoFUS$^{PLD}$ co-condenses with GFP-SS18$^{PLD}$. This movie corresponds to **Fig. 7d**. Scale bar is 10 microns.

**Movie S2:** HEK293T cells co-expressing OptoFUS$^{PLD\text{-}NLS}$ (Cry2-mCherry-FUS$^{PLD\text{-}NLS}$) and GFP-SS18$^{PLD}$. Pre-existing GFP-SS18$^{PLD}$ clusters act as nucleation sites for OptoFUS$^{PLD}$ condensate upon blue light activation. This movie corresponds to **Fig. 7e**. Scale bar is 10 microns.

**Movie S3:** HEK293T cells co-expressing OptoFUS$^{PLD\text{-}NLS}$ (Cry2-mCherry-FUS$^{PLD\text{-}NLS}$) and GFP-SS18$^{PLD}$. Pre-existing GFP-SS18$^{PLD}$ condensates serve as seeds to nucleate OptoFUS$^{PLD\text{-}NLS}$ condensates upon blue light activation. This movie corresponds to **Fig. S18**. Scale bar is 10 microns.
